# Engineered micropillars to unveil oligodendrocyte responses to physical cues

**DOI:** 10.1101/2025.03.16.643578

**Authors:** Eva D. Carvalho, Miguel R. G. Morais, Helena P. Ferreira, Georgia Athanasopoulou, Hendrik Hubbe, Rui Rocha, Maurizio Mattarelli, Beatriz Custódio, Sofia Assis, Shuyao Gu, Kashmyr A. Y. P. Dalang, Samuel L. Rosemore, Wolfgang Losert, María G. Lázaro, Silvia Caponi, Sofia C. Guimarães, Eduardo Mendes, Ana P. Pêgo

## Abstract

Destruction of myelin internodes, oligodendrocyte (OL) apoptosis, and axonal degeneration characterize diseased or aged central nervous systems. While OLs can partially regenerate myelin sheaths, the remyelination process ultimately fails. Tissue mechanical and physical properties, such as stiffness and axonal curvature, play a role in this process. However, the complexity of existing models has hindered studies of OL mechanobiology.

Here, a tissue-engineered model is presented to investigate the impact of stiffness and axonal diameter on OL myelination. The model consists of poly(dimethylsiloxane) micropillars with biologically relevant diameters (1–5 µm), tunable rigidity, and amenable for surface functionalization. The optimized method enables the production of high-aspect-ratio, transparent micropillar arrays, in a reproducible and scalable system, serving as surrogate axons. Additionally, new protocols for quantifying myelin formation are introduced, which can be adapted to any myelination studies.

Softer micropillars accelerate OL differentiation, while rigid ones promote the maintenance of mature OL states. Wrapping of OLs increased with micropillar diameter on rigid substrates, but not on softer ones, suggesting a complex interplay between curvature and rigidity. These processes involve calcium-sensitive channels, histone deacetylases, and microtubules dynamics.

The proposed platform constitutes a versatile and user-friendly system, with applications from fundamental myelin research to drug discovery.

## 1. Introduction

Myelination had a massive impact on the evolution of vertebrates, particularly concerning motor, sensory and high-order cognitive functions of the central nervous system (CNS). The formation of myelin internodes causes insulation of the correspondent axonal segments, so action potential propagation occurs in a saltatory manner, causing an extraordinary increase in the velocity of the nervous impulse propagation ^1^. Oligodendrocytes (OLs) are the responsible cells for myelination in the CNS and their relevance is underlined by the tremendous consequences of their disturbance in several neurodegenerative diseases or even during ageing. Failure of remyelination is linked to axonal degeneration and loss, presumably due to the disruption of the axoglial symbiosis. Consequently, understanding de- and remyelination processes, in all their dimensions, is crucial to characterize tissue degeneration that follows CNS disorders ^2^ and further design new therapies.

OLs inhabit a highly dynamic environment and are continuously subjected to biochemical and biophysical cues that dictate their status: proliferation, migration or terminal differentiation (myelination). External mechanical factors such as matrix stiffness, mechanical strain, macromolecular crowding, and physical confinement have a crucial role in oligodendrocyte precursor cell (OPC) processes’ outgrowth and formation of myelin sheaths ^3^. Acknowledging that no single biochemical signal from the neurons has been identified as a dictator of myelin initiation but that in vivo there is a preference of OLs for ensheathing axons with diameters above a certain threshold (> 0.2 μm diameter) ^4^, the physical features of the axons (caliber and intrinsic curvature) are pointed out as regulators of this process ^5^.

Some studies have shown that in pathological scenarios the mechanical properties of the CNS environment are changed ^6^ but the influence of various mechanical/physical aspects on de- and remyelination remains to be clarified. A critical question is whether these mechanical changes actively drive the disease’s progression or merely serve as correlative indicators.

Some pioneering research studies have advanced our understanding of these phenomena. In vivo studies from Melendez-Vasquez group revealed distinct mechanical profiles between acute and chronic demyelination. While in the acute phase the environmental stiffness decreases due to the loss of a structural component – the myelin, in the chronic phase the mechanical properties are increased, possible due to changes in extracellular matrix (ECM) composition, including the formation of a glial scar ^7^. The same behavior was corroborated by in vitro studies where OLs were cultured in scaffolds with different stiffnesses: cells showed impaired branching and differentiation when cultured in rigid, lesion-like matrices ^8–10^. Interestingly, this phenomenon can also be applied in terms of aging processes: Franklin group demonstrated that the OL ECM stiffens with age, and this is sufficient to cause OL loss of function, a process mediated by Piezo1 (a well-known mechanosensitive ion channel) ^8^. While these studies focused on the effects of stiffness on OL differentiation, the influence of mechanical stiffness on the ultimate function of OLs – myelination, has also started to be revealed. Chew’s group discovered that increased microfiber stiffness inhibits OL myelination, an opposite effect seen for OL differentiation. This effect, regulated through YAP-mediated mechanotransduction signaling, was found to be independent of the fiber diameters tested (1 and 1.5 μm) ^11^. Although OLs exhibit similar myelination profiles within this diameter range, these cells are highly sensitive to the physical properties of axons when the range is expanded. Studies from Chan group using polymeric microfibers as axon surrogates have shown that OLs have increased myelination capacity when cultured on fibers of diameters between 2 – 4 μm, comparing with fibers of 0.2 – 0.4 μm diameter ^12^. Studies from ffrench-Constant lab revealed that OLs not only respond to diameter by varying the number of ensheathments but also by altering their sheath lengths (sheath length was increased in fibers around 2 – 4 μm in comparison with 0.5 – 1 μm) ^13^.

While microfibers have proven reliable for mimicking axonal features and studying myelination, the development of biologically relevant models was significantly advanced by the groundbreaking work of Mei *et al* ^14^. The Chan group pioneered the use of silica cones as a high-throughput system for imaging OL wrapping imaging and screening of different drugs/compounds. This micropillar platform addressed some of the technical challenges associated with microfibers, offering a more straightforward observation of myelin wrapping. Nevertheless, the morphology of these micropillars did not resemble the geometry of biological axons, an issue later resolved by Van Vliet group, who introduced cylindrical micropillars to culture OLs ^15^. These micropillars were prepared from crosslinked resins based on 1,6-hexanediol diacrylate (HDDA) and 4-arm PEG acrylate (starPEG) monomers, presenting lower stiffness compared to silica. Despite this advancement, the reported micropillars had diameters of 10 and 20 μm which largely exceeded the typical average CNS axonal diameters of 1-2 μm. More recently, the same group refined micropillars fabrication methodology and were able to reduce the structures’ diameters to 5 μm ^16^.

All these previous models (for a summarized overview of the above referenced reports see Table S1 in SI) are, to varying degrees, technically challenging to implement in a standard biology laboratory. Furthermore, to the best of our knowledge, the relationship between axon diameter and stiffness has not yet been systematically examined for an extended biologically relevant range of axonal diameters.

Here we propose a new simple system to study the influence of substrate stiffness and axonal structure on OLs behavior. Our platform consists of an array of poly(dimethylsiloxane) (PDMS) micropillar structures with biologically relevant dimensions (ranging from 1 to 5 μm diameter and 10 μm length). These micropillars have tunable stiffness and their transparency facilitates straightforward visualization of live OL interactions with the micropillars. The simplicity of PDMS fabrication ensures that the system is easily reproducible and accessible for implementation in standard laboratory settings. Overall, these features make them well suited for adaptation to a high-throughput testing system.

Exploring this versatile setup, we show that softer PDMS substrates accelerate early differentiation, and that thicker micropillars stimulate OL wrapping. For the first time, we demonstrate the involvement of calcium sensitive channels (transient receptor potential cation channels, TRPV1 and TRPV4) and histone deacetylases (HDAC3) in response to stiffness variations. We also reveal that microtubules dynamics are potential mediators of OL response to different curvature structures. Finally, we present a simple workflow for automated image quantification of myelin formation around the micropillars, which can be easily applied in any related study.

## 2. Results

### 2.1. Development of tunable PDMS micropillars

We engineered transparent, mechanically tunable PDMS axonal surrogates with biologically relevant dimensions (ranging from 1 to 5 μm in diameter, comparable to in vivo CNS myelinated axons ^4^) and a high-aspect ratio, mimicking axonal features such as caliber, morphology and curvature. Micropillars were produced through a replica molding technique, peeled off in the presence of isopropanol and dried by critical point drying (Figure 1A and Figure S1). The structures presented the desired cylindrical shape and dimensions comparable with the projected theoretical ones (Figure 1B and Table S2).

**Figure 1.**
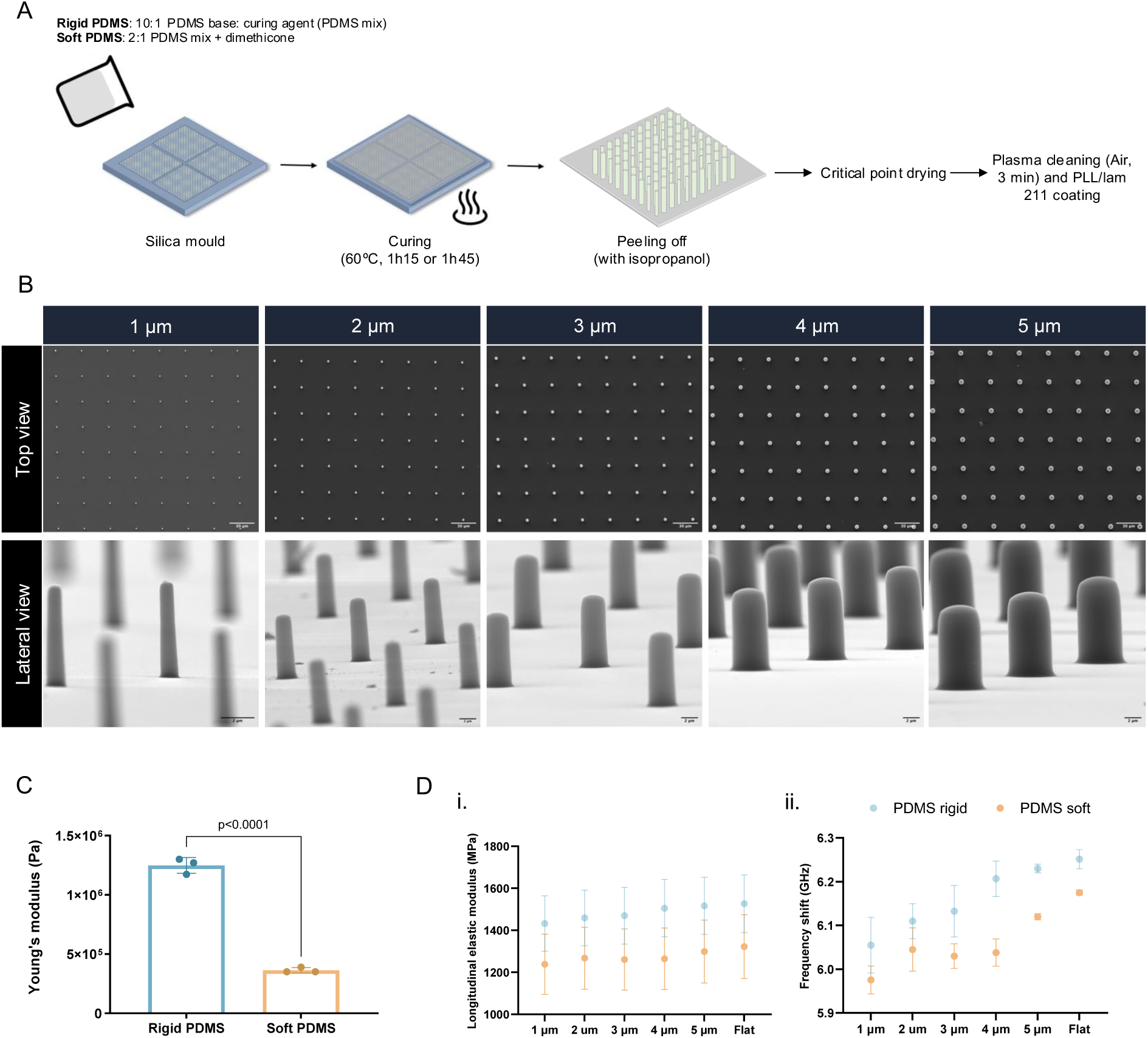
Poly(dimethylsiloxane) (PDMS) micropillars resemble axonal physical structure. **(A)** Micropillars were produced by casting PDMS prepolymer into a silica wafer mold followed by curing at 60 °C for 1 h 15 min or 1 h 45 min (rigid and soft micropillars, respectively). For rigid PDMS, a 10:1 base-to-curing agent ratio was used and for soft PDMS dimethicone was added to the mix (2:1). Micropillars were peeled off in the presence of isopropanol, followed by critical point drying to avoid structural collapse and plasma treatment. A coating of poly(L-lysine) and laminin 211 was used to mediate cell adhesion. **(B)** Morphological characterization of micropillar arrays through scanning electron microscopy (SEM). Micropillars ranging from 1 to 5 μm diameter were produced, with a theoretical height of 10 μm and an inter-distance between micropillars of 30 μm. Scale bar is depicted in the images (top view 30 μm, lateral view 2 μm). **(C)** Young’s modulus from rigid and soft PDMS formulations. Values were obtained by rheological measurements on PDMS 4 mm diameter discs. n=3 PDMS discs per condition, for each type of oscillation test. Presented values correspond to mean ± SD. Statistical analysis was performed using One-way ANOVA (p<0.05). **(D)** Longitudinal elastic modulus (i) and frequency shift of PDMS micropillars (ii) measured by Brillouin microscopy. Results show mean ± SD (n>3 micropillars per condition).

We significantly reduced the PDMS Young’s modulus (Figure 1C) by introducing a low-viscosity PDMS dimethicone, corresponding to a significant advance in the fabrication of elastomeric structures with a great aspect ratio. To understand if different micropillars’ diameters have different rigidities, we used Brillouin microscopy, a contactless and non-destructive technique to assess mechanical properties. The results confirmed an overall difference in the stiffness of the two different PDMS formulations, with the stiffer one exhibiting a longitudinal modulus of around 1.5 GPa and the softer around 1.3 GPa (Figure 1D). In addition, we found no differences in the longitudinal modulus among different diameter micropillars, which further allowed us to independently control stiffness and diameter effects on cells.

In addition, we prove the amenability of the proposed structures to be coated by biologically relevant proteins, following plasma treatment. As can be seen in Figure S3, PDMS micropillars were homogeneously coated with laminin 211, a facilitator of OPC adhesion and differentiation^17^.

It is also important to highlight that our methodology enables the fabrication of a diverse array of micropillar platforms (Figure S1), ranging from single-diameter to mixed-diameter configurations, being the later a closer approximation to the natural OL environment.

Overall, we report a new method for producing high-aspect ratio, transparent micropillar structures in an easy, rapid, reproducible, low-cost, and scalable way, that can be easily implemented in any standard biology lab.

### 2.2. Oligodendrocyte differentiation and myelin wrapping is recapitulated on micropillars

We next defined the optimal OL culture conditions on micropillars (Figure S3), including cellular density, and evaluated key cell-related parameters: survival, metabolic activity, and differentiation. We found the structures to be cytocompatible with cellular metabolic activity not being affected by the biomaterial structure (Figure S4). We observed that when seeded on the platforms OPCs are highly dynamic in the first days of the culture, closely interacting with multiple micropillars until establishing stable contacts at day 5 of differentiation (D5 DIFF, Movie S1-2, Figure S5). OPCs were not only able to differentiate in myelin-forming OLs but also wrap the micropillars (Figures 2A and 2B) while displaying the typical OPC/OL genetic markers across every structure tested (rigid *vs* soft PDMS platforms) (Figure 2C). Proteolipid protein (PLP1) was among the highest expressed marker, which is in line with the high abundancy of this protein in the CNS myelin ^1^. Similarly to what happens in vivo, we also found that individual OLs were able to wrap around multiple micropillars and that the wrapping process occurred not only radially but also longitudinally, along the pillars’ height. Notably, at later time points of differentiation, we observed that cells were able to produce more than one myelin concentric layer around the pillars (Figure 2D), in accordance with other published works with similar in vitro platforms ^13,14^. In addition, OLs presented calcium activity, a strong indicator of their functionality (Figures 2E and 2F, Figure S6 and Movies S3A-C).

**Figure 2.**
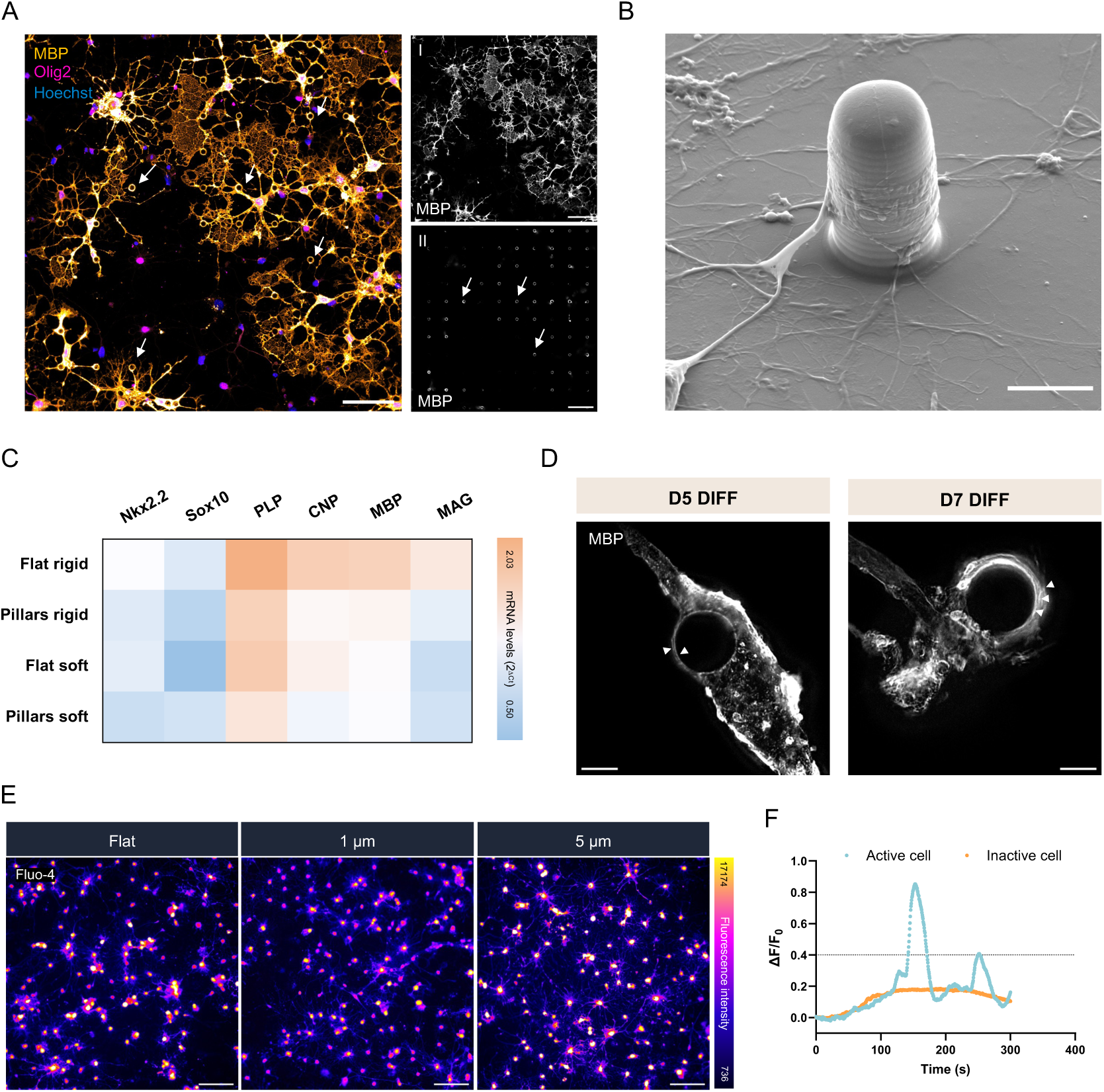
Oligodendrocytes (OLs) differentiate, myelinate and are functionally active when cultured on micropillar structures. **(A)** Representative confocal microscopy images of OLs on micropillars at day 5 (D5) of differentiation (DIFF) expressing myelin basic protein (MBP, orange) and Olig2 (magenta) (nuclei counterstained with Hoechst, blue). **I** and **II** represent bottom and top z-planes of OLs on micropillars, respectively (MBP, white). OLs extend long myelin sheaths and wrap around the micropillars as depicted by the myelin rings observed in II and depicted by the arrows. Scale bar indicates 50 μm. **(B)** Scanning electron microscopy (SEM) of an OL wrapping around a micropillar at D5 DIFF. OLs wrap micropillars axially but also longitudinally. Scale bar indicates 5 μm. **(C)** mRNA levels of OL specific genes on micropillars and flat PDMS surfaces at D5 DIFF. Levels were normalized to the *Oaz1* housekeeping gene, and results indicate mean values from 3 independent experiments (8 mixed diameter micropillar-/flat-structures pooled together). At this differentiation stage, OLs expressed their common markers, highlighting the feasibility of the PDMS structures in promoting OL lineage commitment and differentiation patterns. **(D)** Stimulated emission depletion microscopy (STED) representative images of OLs wrapping the micropillars and expressing MBP (white). Some micropillars displayed thick myelin sheaths with more than one visible concentric layer (noted by the arrows). Scale bar indicates 2 μm. **(E)** Representative fluorescent images of OLs loaded with the calcium indicator Fluo-4 at D5 DIFF. The image pseudo colours indicate differences in pixel intensities. Scale bar indicates 100 μm. **(F)** Representative calcium signals overtime of an active (blue) and inactive (orange) cell at D5 DIFF. Calcium events were scored as increases of ΔF/F_0_ equal to or more than 40% of the baseline.

We and others have previously reported the feasibility of platforms such as microfibers and micropillars for supporting OL differentiation and wrapping ^13–15,18–21^. Our new PDMS-based micropillar model is well-suited for imaging techniques, enabling easy cell tracking, and visualization of myelin membrane formation.

### 2.3. Micropillar diameter and stiffness impact OL differentiation and wrapping capability

In myelin research, a long-standing question remains: why do some axons get myelinated while others, even in regions of abundant white matter, never develop myelin sheaths? The current hypotheses are centered around the inherent capability of OLs to produce myelin sheaths, contingent upon the presence of appropriate physical cues for this process. Besides the axon’s diameter, other mechanical properties are expected to impact myelination. Exploring our platform, we next investigated if OL differentiation and wrapping capacity was modulated by the different physical characteristics of our PDMS micropillar structures, namely diameter and rigidity. We developed a rapid workflow to measure myelin wrapping on axon-mimicking structures, involving image acquisition followed by analysis using open-access software, making it easy to implement in any conventional laboratory (Figure 3A). By imaging the top z-plane of fluorescently labeled OL images featuring myelin markers, one can identify, segment, and correlate the myelin rings with the micropillars structures. From these images (Figure 3B) different outputs can be taken, including (1) the branching ability of the cells (based on the staining of the F-actin marker); (2) the number of myelin basic protein (MBP) and Olig2 positive cells; (3) the area occupied by the myelin marker MBP; (4) the intensity of the MBP either in overall measurements or in the micropillar region; (5) and the number of wrapped micropillars.

**Figure 3.**
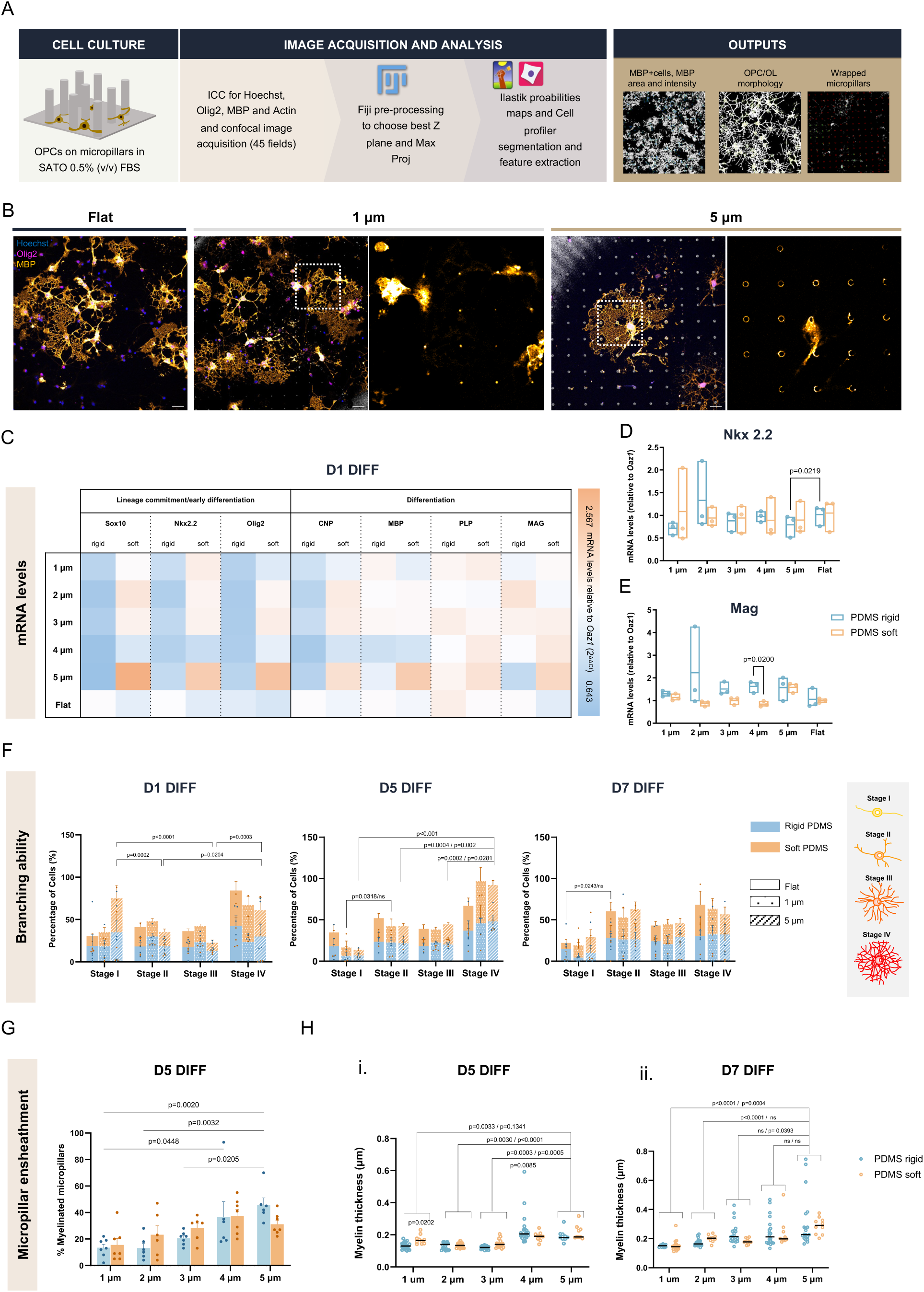
Oligodendrocyte (OL) differentiation and wrapping ability is dependent on micropillar diameter and rigidity. **(A)** Diagram of the sequential steps for image acquisition and analysis. OLs were cultured on PLL/lam211 coated micropillars or flat surfaces iand fixed at D1, 5 or 7 DIFF. Confocal images were acquired for myelin basic protein (MBP, orange) and Olig2 (magenta) markers, while nuclei were counterstained with Hoechst (blue). **(B)** Representative images of OLs cultured on flat, 1 μm and 5 μm micropillar platforms. Scale bar indicates 25 μm. Zoomed in images show regions of wrapped micropillars. **(C)** mRNA levels of OLs cultured on different diameter and stiffness micropillars at D1 DIFF. Lineage commitment (Sox10, Nkx2.2, and Olig2) as well as differentiation (CNP, MBP, PLP and MAG) markers were analyzed. Results show mean values of levels normalized to Oaz1 housekeeping gene and to respective flat surfaces (2^-ΔΔCt^, n>3 independent experiments, 8 micropillar platforms pooled together). Statistical analysis performed using a two-way ANOVA. **(D)** and **(E)** Nkx2.2 and Mag mRNA levels of OLs cultured on different diameter and stiffness micropillars at D5 DIFF. Results show mean values of levels normalized to Oaz1 housekeeping gene and to respective flat surfaces (2^-ΔΔCt^, n>3 independent experiments, 8 micropillar platforms pooled together). Statistical analysis performed using a two-way ANOVA. **(F)** OL branching ability at D1, 5 and 7 DIFF. Quantification was performed based on the “Collar Occupancy method” ^22^. Results represent mean ± standard deviation (SD), n > 5 micropillar platforms from 4 independent experiments. Two-way ANOVA. **(G)** Percentage of wrapped micropillars by OLs in different diameter and stiffness micropillars. Results show mean ± SD, n>6 micropillar platforms analyzed from 4 different independent experiments. Two-way ANOVA. **(H)** Myelin thickness of OLs on micropillars at D5 (i) and D7 DIFF (ii). Measurements were performed from images acquired by STED microscopy. Six measurements per z-plane from a total of four planes per micropillar were made. At least 10 micropillars per condition were acquired. Real values of the diameter of the micropillars were estimated based on the length of the mid-plane acquired. Results show median values. Statistical analysis was performed using Two-way ANOVA.

Generally, we observed a tendency for the increased expression of lineage commitment/early differentiation markers at D1 DIFF for softer micropillars in comparison with rigid ones (Figure 3C), an effect that was not visible at later time points of culture (Figure S7). Of note, *Nkx2.2* expression was significantly decreased for 5 μm rigid micropillars in comparison with the corresponding flat surfaces at D5 DIFF (Figure 3D). This suggests a potential arrest of the precursor state in cells that are in contact with flat 2D substrates. Additionally, at the same time point, we detected a general increase in the genetic expression levels of *Mag* on rigid substrates in comparison with softer ones, exhibiting statistical significance particularly for the 4 μm micropillars (Figure 3E). At D7 DIFF the same trend was seen but for the *Mbp* gene (Figure S7), indicating that while softer structures might accelerate the early differentiation phase, rigid substrates promote the expression of genes associated with OL differentiation and maturation. However, the effect of the stiffness was not evident in terms of cellular morphology and micropillar wrapping capacity (Figures 3F and 3H, S8A). Interestingly, we found that when cells are in contact with stiffer substrates, the effect of micropillar diameter is more pronounced than when these are cultured on softer environments. This was substantiated by the number of myelinated micropillars, which was progressively higher from 1 to 5 μm micropillars only on stiffer substrates (Figures 3G and S8C). When assessing the MBP thickness around micropillars using stimulated emission depletion microscopy (STED), we observed that this parameter generally increased as a function of the micropillar diameter at D5 and 7 DIFF (Figure 3H). In vivo, it is reported that myelin thickness values vary proportionally with axonal fiber diameter, further emphasizing the biological relevance of our model ^4^. In addition, we also recorded that the occupied area by MBP in the micropillar region decreased from D5 to D7, an indicator of more condensed myelin regions (Figures S8D and S8E).

Overall, we observed that OLs’ exhibit better performance in terms of differentiation on micropillars than on flat surfaces. Firstly, on flat surfaces the OPC/OL population distribution across different stages of differentiation is evener, whereas on micropillars most cells are in more mature stages (Figure 3F). Then, OLs on micropillars occupy, in general, higher areas than on flat surfaces (Figure S8B).

All together our results indicate a strong impact of physical cues on the capacity of myelin wrapping by OLs, with larger and the softer micropillars being favored for these processes. Furthermore, the relationship between diameter and stiffness was found to be close and myelination intricately dependent on this interconnection.

### 2.4. OLs perceive micropillar cues through calcium-sensitive channels, impacting epigenetic regulators and microtubules dynamics

We next sought for potential biochemical candidates that regulate the interaction of OLs with micropillars of varying stiffness and diameter. First, we analyzed the gene expression levels of cytoskeletal proteins, mechanosensitive channels, transcriptional activators, and epigenetic regulators involved in mechanosensing mechanisms (Figure S9A).

Integrins are membrane-associated receptor proteins highly involved in the process of cell-ECM interaction, being the subunit α6β1 a mediator of the contact between OLs and neurons, and also a sensor of the axonal size ^23^. Signaling through integrin receptors can occur via focal adhesion kinase (FAK), proto-oncogene tyrosine-protein kinase Src and phosphoinositide 3-kinase (PI3K) pathways, with Ras homolog gene family member A (RhoA) being one of the major mediators of this process with extensive effects on cellular function. RhoA is implied in OL differentiation and myelination ^24^. We observed relatively stable levels of the expression of the integrin receptor *Itgb1* as well as the *RhoA* genes (Figures S9B and S9C). Yet we noted a tendential progressive increasing of the expression of these genes as a function of culture time for the rigid micropillars. This could indicate increased adhesion capacity of OLs to these substrates as well as enhanced cytoskeletal tension and signaling driven by substrate rigidity. Mechanical stimuli are also perceived through mechanosensitive channels, as the case of the ones from the TRPV family. We observed a consistent enhancement of the expression of *Trpv1* and *Trpv4* for softer structures in comparison with rigid ones (Figures 4A and B). Although still scarce, literature reports that *Trpv4* channels increase OPC proliferation ^25^. Interestingly, the effect of the expression of *Trpv4* decreased as function of micropillar diameter, suggesting a differentially regulated mechanism of mechanosensing occurring in the presence of larger diameter pillars.

**Figure 4.**
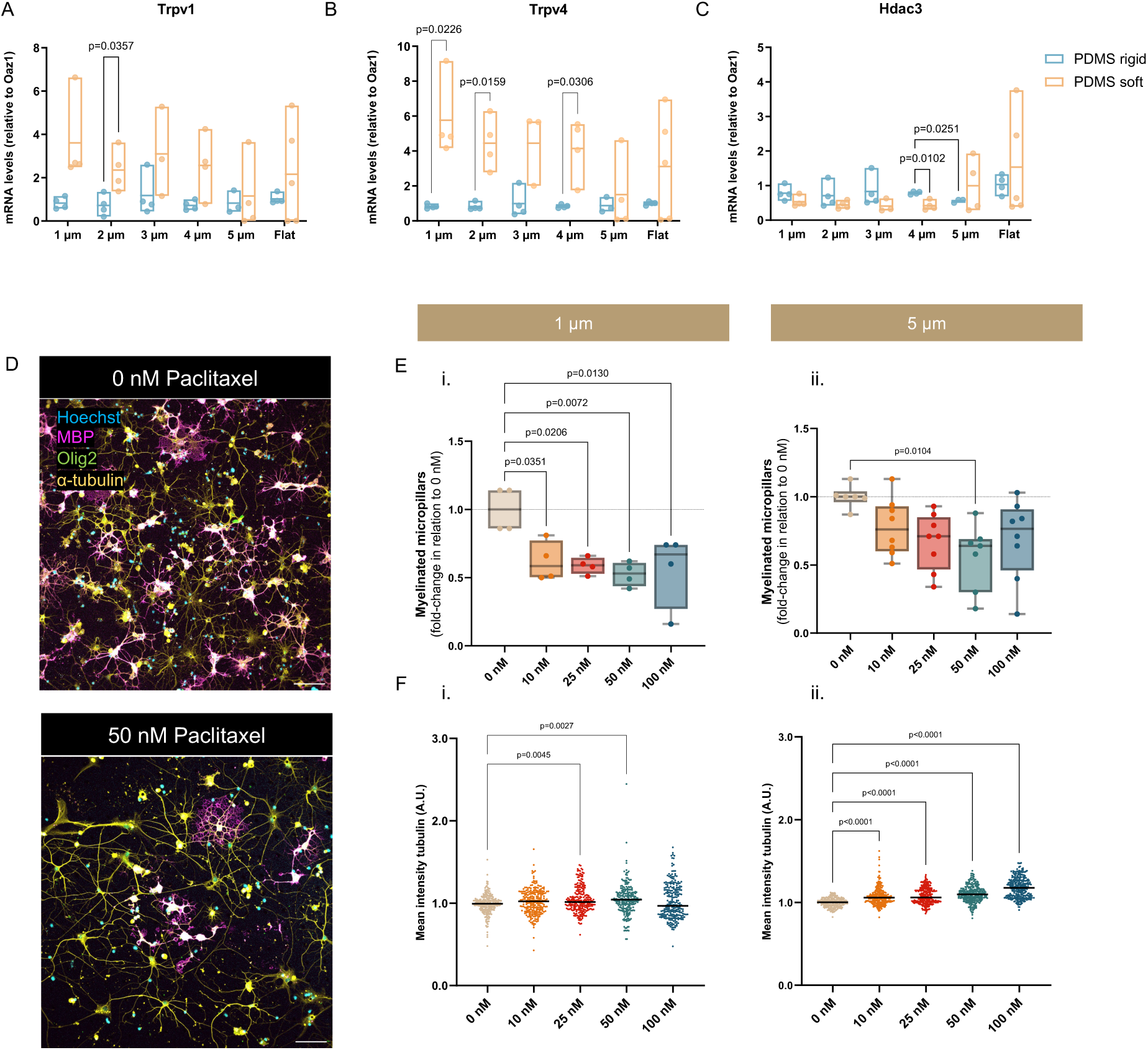
Oligodendrocytes (OLs) respond to micropillars’ mechanical cues through differentially expression of calcium sensitive channels and epigenetic modulators, as well as through cytoskeletal alterations. mRNA expression levels of *Trpv1* (A), *Trpv4* (B) and *Hdac3* (C) of OLs cultured on micropillars at day 1 of differentiation. Results show mean values, n>3 independent experiments (eight micropillar platforms pooled together for each condition per independent experiment). Statistical analysis was performed using Two-way ANOVA. **(D)** Representative images of OLs cultured in the presence of the microtubule stabilizer Paclitaxel. **(E)** OLs were cultured in the presence of Paclitaxel for 12 h and a reduction of the number of myelinated micropillars was observed both in platforms with 1 μm (i) or 5 μm (ii) micropillars. Values show mean ± SD. Statistical analysis was performed using One-way ANOVA. **(F)** Quantification of the alpha-tubulin area of OLs after treatment with Paclitaxel on 1 μm (i) or 5 μm (ii) micropillars.

We also investigated the expression of the most commonly studied mechanosensing transcription factors, YAP and TAZ (Figure S9D). Their activity is regulated by the conformation and tension of the F-actin cytoskeleton, which, in turn, depends on the characteristics of the substrate to which the cells adhere ^26^. Although not statistically significant, a tendentially augmented expression levels of *Yap* and *Taz* was noticed for OLs cultured on softer structures, being this more pronounced on the cells cultured in the presence of the smaller diameter micropillars. *Yap* and *Taz* are normally activated and translocated to the nucleus when cells are in contact with stiffer substrates ^26^. Assuming *Yap/Taz* increased activation on stiffer structures, cells may not need to continuously upregulate mRNA levels as actively as they need on softer environments.

Concomitantly with transcriptional changes, some studies also link the effect of external forces with epigenetic alterations in OLs ^27,28^. We looked for differences on the epigenetic regulators of OLs cultured on the microstructures, particularly the expression of *Hdac* encoding genes. HDACs are enzymes known to regulate gene expression by modifying the structure of chromatin through removal of acetyl groups from lysine residues on histone proteins ^29^. Here, we observed a decreased expression of *Hdac2* and *Hdac3* (Figure 4C, S9D) for OLs growing on softer structures in comparison with stiffer ones. The tendency occurred for all micropillar diameters tested with the exception of the 5 μm size, where the expression levels were lower for the rigid structure and significantly different when comparing with the 4 μm. Decreased expression of *Hdac2* and *3* may lead to a more open chromatin structure allowing a broader range of genes to be transcribed. This can imply that that OLs are more flexible to adapt to various cues on softer substrates in relation to stiffer ones, whose restrictive chromatin structure might promote higher levels of differentiation and OL specialization ^30^.

Overall, the mRNA levels of the targeted genes analyzed corroborates an interdependence between axonal caliber and stiffness in the regulation of OL differentiation and wrapping.

Acknowledging that OLs wrap more efficiently micropillars of large diameter, we next questioned whether interfering with the sensing of different curvatures would alter the wrapping process by OLs. Our hypothesis is that by reducing the flexibility of microtubules, micropillar ensheathment would be hampered. The high curvature of 1 μm micropillars can pose a mechanical challenge, requiring highly flexible microtubules to accommodate the tight bends. In contrast, micropillars with larger diameters (5 μm) provide gentler curvatures, which may allow microtubules to bend more easily. By stabilizing the microtubules with Paclitaxel (Taxol®) we expected that the capacity of the OL membrane to bend in the presence of micropillars would decrease. As it stabilizes microtubules, Paclitaxel could also lead to increased intracellular stiffness, which in turn could impair membrane wrapping. A reduced number of wrapped 1 and 5 μm micropillars was observed when OLs were cultured in the presence of 50 nM of Paclitaxel for 12 h (Figures 4D and E), whereas the intensity levels of the tubulin progressively increased in a dose dependent manner (Figure 4F). These differences might be specifically linked to changes in cytoskeleton dynamics as alterations on cell viability and MBP positive cells were not observed (Figure S10). The response of OLs to Paclitaxel was more pronounced when OLs were cultured on 1 μm micropillars, highlighting that, increased curvatures impact the process of wrapping to a larger extent.

These findings underline the close crosstalk between epigenetic modifications, differentially expressed mechanosensitive channels and cytoskeleton dynamics that orchestrate the optimal myelin ensheathment (Figure 5).

**Figure 5.**
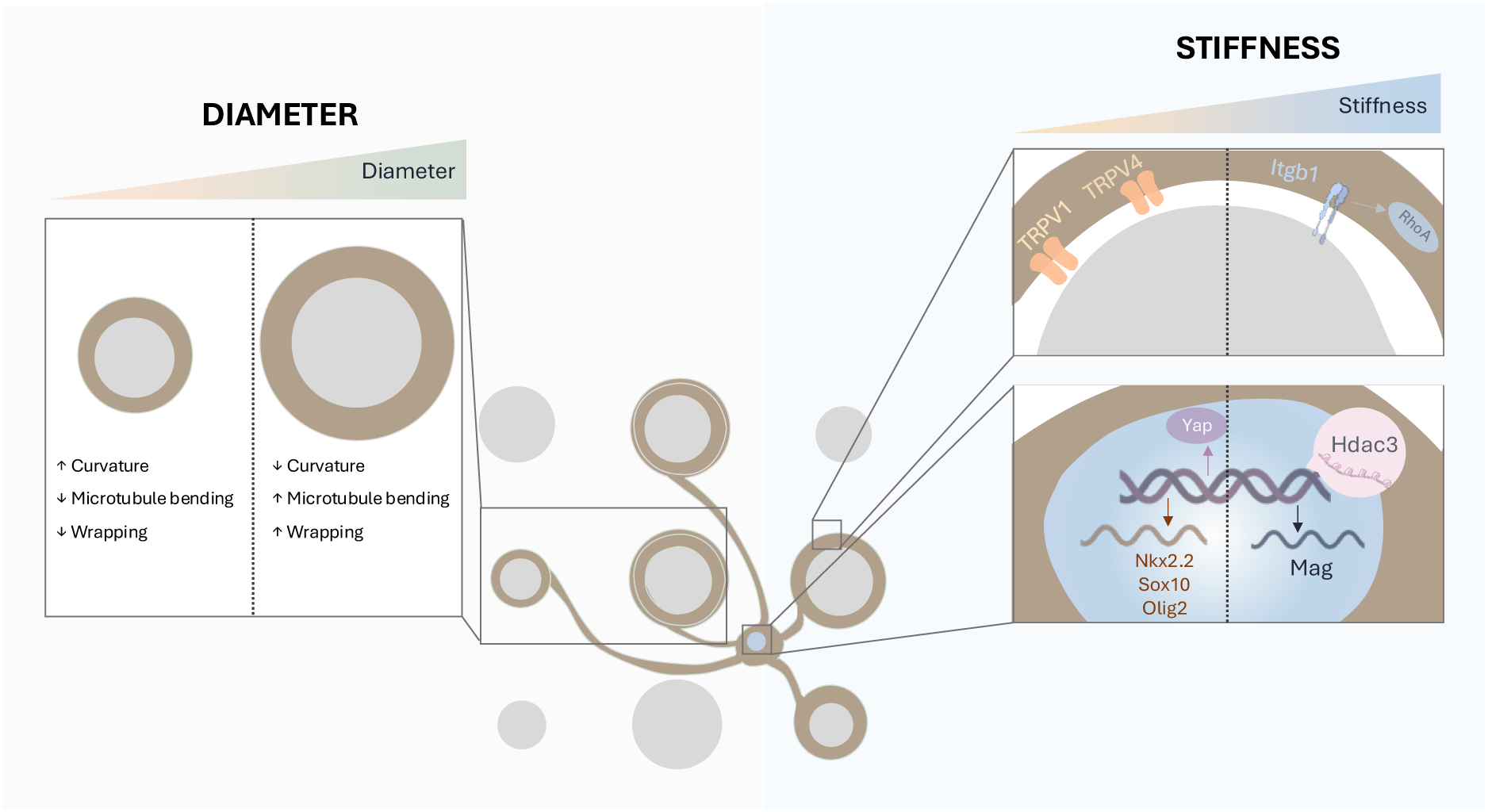
Proposed model for oligodendroglial sensing of micropillar rigidity and diameter. Early differentiation is accelerated on lower mechanical tension substrates while more sustained differentiation and maturation occurs in stiff substrates. Increased integrin and RhoA levels could enhance adhesion capacity of OLs as well as activation of cytoskeletal tension signaling pathways. In addition, *Hdac3* increased expression contribute to a cellular state that favors stability, an important feature required for myelination. On the contrary, on softer substrates, lower mechanical tension might be more permissive for initial differentiation signals to emerge, favoring gene expression and cellular changes characteristic of early differentiation without fully commitment to a mature state. Diameter also dictates OL fate: large diameters and therefore low curvatures are more permissive for wrapping, with microtubules bending easily. On the other hand, smaller diameters have high curvatures which hampers microtubule binding and consequently wrapping.

## 4. Experimental Section/Methods

### Design and production of silica wafers

The PDMS micropillar structures were created using silicon wafer molds. Molds were produced using electron-beam lithography and deep reactive ion etching (DRIE). The surface of a silicon wafer was primed to enhance resist adhesion by spin-coating with AR 300-80 (Allresist GmbH Germany) at 4000 rotations per minute (rpm) for 1 min and baked subsequently at 180 °C for 2 min. Next, the wafer was spin-coated with positive resist AR-P6200.18 (Allresist GmbH Germany) at 4000 rpm for 1 min and soft baked at 180 °C for 3 min. To avoid cracks in the structured resist, the cooling down after soft bake was gradually performed on the hotplate to reduce stress (ca. 10 min to 90 °C). Electron-beam writing was done using a dose of 300 µC cm^-2^ (Leica EBPG 5000+ 100 kV), followed by bath development in pentyl acetate for 5 min, 4-Methylpentan-2-one (MIBK)/2-propanol (1:1 v/v) 3 min and 2-propanol 3 min. For the etching process, a stepped Bosch process DRIE recipe cycle was used. Passivation was applied for 2.8 s using C_4_F_8_ (300 SCCM) at 1500 W ICP power, etching continued using SF_6_ for 2.2 s (200 SCCM) with higher ICP power 1800 W and additional 6 W LF power, followed by the main isotropic etching with SF_6_ for 2.6 s (300 SCCM) at ICP power 1800 W. After plasma surface activation (40 mbar, 100 W, 3 min), the wafer molds were exposed to fumes of 200 µl trichloro(1H,1H,2H,2H-perfluorooctyl)silane (CAS 78560-45-9) in a desiccator under vacuum for 4 h and are then ready for PDMS molding. The molds consisted of 7×7 mm squares with cylindrical cavities of 10 μm depth and different diameters (1, 2, 3, 4 and 5 μm), distanced 30 μm from each other. Different designs were produced (Figure S1).

### Fabrication of polydimethylsiloxane (PDMS) micropillars’ platforms

Polydimethylsiloxane (PDMS) was obtained from Silastic® T-2 Translucent Base (viscosity 50,000 mPa·s) and Curing Agent (Dow Corning). PDMS prepolymer was prepared in a 10:1 base-to-curing agent ratio. Micropillars were fabricated by casting PDMS prepolymer into a silica wafer mold and curing at 60 °C for 1 h 15 min. After polymerization, the micropillars were peeled off from the mold and cut out, immersed in 2-propanol. PDMS flat surfaces (used as controls) were fabricated similarly.

To decrease the substrate stiffness, micropillars were also manufactured from a mixture of PDMS (10:1 ratio) with 360 Medical Fluid (viscosity 100 cSt, Dow Corning) (designated dimethicone) and mixed in a 2:1 ratio total PDMS-to-dimethicone ratios (w/w). Polymerization was carried on for 1 h 45 min.

PDMS micropillars were washed with an increasing gradient of ethanol:2-propanol (0:100, 50:50, 100:0) for a complete substitution of 2-propanol for ethanol and were critical point dried (Polaron Range CPD7501, Quorum Technologies). Critical point drying was performed with 7 cycles of flush (at 4 °C, 800 psi) to substitute ethanol for liquid CO_2_. PDMS platforms were stored in a desiccator until further use.

### Scanning electron microscopy (SEM)

Samples were coated with a gold/palladium thin film by sputtering, using the SPI Module Sputter Coater equipment. Coating was initially performed with a 45° tilt for 40 s, and then from a top view for 40 s.

Samples were then observed using the High-Resolution Quanta 400 FEG ESEM, from a top view (no inclination) and lateral view (90° tilt). The diameter at the mid-length, at the base, and at the top of the micropillars, its length, and distance between them were estimated in different micropillars. Measurements were performed in three images per different condition.

For cellular studies, OLs were fixed in glutaraldehyde (2.5% (wt/v) diluted in cacodylate buffer 0.1 M pH 7.2-7.4 (prepared in phosphate buffered saline, PBS, without calcium or magnesium) for 30 min at room temperature (RT) under gentle stirring (50 rpm). Samples were washed with cacodylate buffer twice and then dehydrated in serial diluted ethanol solutions of 50, 60, 70, 80, 90, and 99% (v/v) for 10 min in each dilution. Afterwards, samples were dried in hexamethyldisilazane (HMDS) and coated with gold/palladium for 80 s (from the top).

### Rheology

The viscoelastic properties of the micropillars were characterized by rheology (Kinexus Rheometer, Malvern Instruments Ltd.). Four mm-diameter flat discs (cut with a puncher) of PDMS were used to perform oscillation tests with a 4 mm parallel-plate geometry, at 37 °C and 0.1-0.2% compression. To determine the linear viscoelastic region (LVR), 3 amplitude sweep tests were performed at a constant oscillation frequency of 1 Hz. Once the LVR for each material was defined, three oscillation frequency sweep tests were conducted. For amplitude sweep tests, samples were submitted to strains in the range of 0.5-150%, at a constant frequency of 1 Hz (10 points per decade). For frequency sweep tests, samples were submitted to oscillations in the range of 0.1-10 Hz, at a constant strain of 5% (10 points per decade). Values of G*, G’ and G” were determined from both types of tests. For each type of oscillation test, n=3 technical replicates were analyzed. The tolerance range of deviation for G*, G’ and G” around the plateau value was ± 5%. Stiffness of the materials used in micropillars, as defined by the Young’s modulus (E), was then calculated using G*. For linear isotropic materials, G*, and E are related through the equation *E* = 2*G* ∗ (1 + *v*), where *v* stands for the Poisson’s ratio ^31^. The PDMS 4 mm-diameter discs were admitted to be isotropic (*i.e.* mechanical properties have identical values in all directions) and *v* = 0.5 was used as a reference value from literature for the Poisson’s ratio ^32^.

### Brillouin microscopy

PDMS micropillars were characterized in terms of longitudinal modulus (M) by Brillouin microscopy. The Brillouin spectra are acquired using the Tandem Fabry Perot interferometer (TFP-2 HC). In brief, the light of a single mode diode-pumped-solid state Spectra-Physics Excelsior laser operating at λ=532 nm is focused on the pillars using a 20X infinity-corrected apochromatic microscopic objective (Mitutoyo M-Plan Apo 20×) with a very long working distance of 20 mm, and numerical aperture of 0.42. The Brillouin peaks are fitted using a DHO function

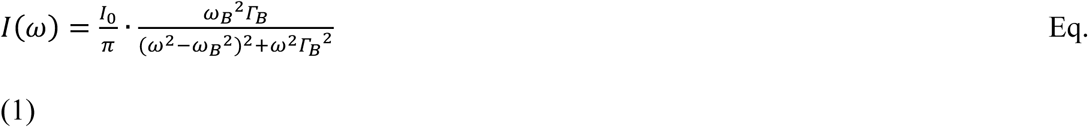

which provides the frequency position of Brillouin peak, ω_B_ and its width, Γ. The longitudinal elastic modulus can be obtained by the Brillouin frequency shift as

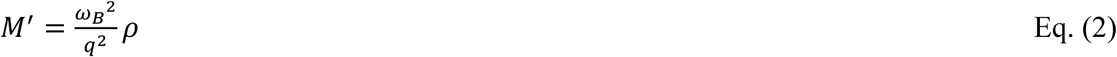

where q in the backscattering configuration is q= 4πn/λ, being λ the wavelength of laser light, n is the refractive index and ρ the density of the investigated material.

The density of the different PDMS structures was estimated using a 100 mL capacity pycnometer. Briefly, the mass of the pycnometer was measured (*m_pyc_*) and then seven micropillar arrays previously critical point dried were introduced in the pycnometer and the total mass weighed again (*m_pyc_*_+*pillars*_). Arrays were afterwards removed from the pycnometer and plasma treated, as described before. The pycnometer was filled with deionized water and its mass measured (*m_pyc_*_+*H2O*_) followed by weighing the pycnometer filled with water and micropillars (*m_pyc_*_+H_2_O+*pillars*_). The volume of the water with (*v*_H_2_O+*pillars*_) or without pillars (*v*_H_2_O-*pillars*_) were estimated based on the water density (0.997 kg/m^3^ at 25 °C) using the following equation:

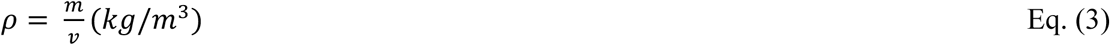

where ρ is the density, *m* is the mass and *v* the volume.

The volume of the micropillars (*v_pillars_*) was estimated based on the difference of the volume of water with and without micropillars. The mass of the micropillars (*m_pillars_*) was calculated as the difference of the *m_pyc+pillars_* and *m_pyc_*. The density of the micropillars was then estimated based on Eq. 1. Measurements were conducted at room temperature (approximately 22/23 °C).

The obtained values were:

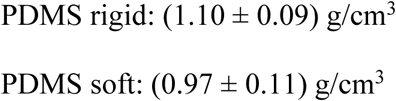

The refractive index was determined using an Abbe refractometer (DR-A1). PDMS films of 25×8×3 mm were produced (seven per condition) and one measurement per condition was performed. Recordings were conducted between 19.5-19.7 °C. The obtained values of refractive index measured at λ = 589 nm were:

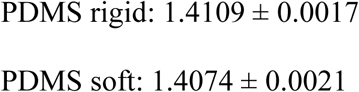

### Mixed glial cell (MGC) cultures

All procedures involving animals and their care were performed in agreement with institutional ethical guidelines (IBMC/INEB/i3S), the EU directive (2010/63/EU) and Portuguese law (DL 113/2013). Experiments described here had the approval of Portuguese Veterinary Authorities. Animals had free access to food and water, being kept under a 12 h light/12 h dark cycle.

Primary cultures of OPCs were obtained as previously described ^18,33,34^. Briefly, mixed glial cells (MGC) cultures were isolated from the brains of post-natal day 0 to day 2 (P0-P2) Wistar Han rats. Pups were sacrificed by decapitation and brain removed and dissected in Hank’s Balanced Salt Solution without calcium and magnesium (HBSS, Sigma) supplemented with 2% (v/v) penicillin/streptavidin (P/S) on ice. Cortices were isolated by removal of the meningeal tissue, the cerebellum, the basal ganglia and the hippocampus. Isolated cortices were then digested in HBSS without calcium or magnesium supplemented with trypsin (0.0025% (wt/v)) and 0.001 mg/mL DNAse I (Applichem LifeSciences) for 15 minutes at 37 °C. Dissociated cortices were cultured in 10 µg/mL poly(D-lysine) (PDL, Mw >300,000 Da, Merck A003E) coated 75 cm^2^ flasks (30 min, 37 °C, and 5% CO_2_) and maintained in Dulbecco’s Modified Eagle Medium (DMEM) supplemented with 10% (v/v) heat-inactivated (30 min, 56 °C) foetal bovine serum (FBS, Sigma) and 1% (v/v) P/S. Cells were grown in a cell culture incubator at 37 °C and 5% CO_2_. When confluence was reached (∼12 days) the flasks were submitted to a pre-shake (200 rpm, 2 h, 37°C) causing detachment of microglia. Medium with detached microglia was removed and culture medium was refreshed. Following a 2 h incubation period (37°C, 5% CO_2_), MGCs were submitted to shake-off (220 rpm, overnight, 37 °C), resulting in detachment of OPCs and remaining microglia (while astrocytes remain adhered during this procedure). Medium with detached OPCs and microglia was transferred to non-coated and non-treated Petri dishes (90 mm in diameter) and incubated for 2 h (37 °C, 5% CO_2_). Microglia adheres faster to non-treated substrates than OPCs, allowing for the removal of the remaining microglia. Suspension medium containing OPCs was passed through a 40 μm nylon cell strainer (Falcon) to remove astrocyte clusters that could possibly have detached during shaking and centrifuged for 10 min, 500 x g, at RT. Cells were then resuspended, and live cells counted using the Trypan blue assay.

### Oligodendrocytes cultures on PDMS micropillars

To increase PDMS platforms hydrophilicity, promote exposure of reactive groups and sterilization, they were plasma treated using the Diener Electronic Zepto Plasma Cleaner, at a low pressure (0.8 mbar), 30% intensity for 3 min. Coating of PDMS surfaces with 50 μg/mL poly(L-lysine) (PLL, Mw 30,000-70,000, Sigma-Aldrich P2636) was performed immediately after plasma treatment (overnight, 37 °C, 5% CO_2_). After washing twice with sterilized type II water, PDMS platforms were coated with laminin 211 (Biolamina LN211, 10 μg/mL) and incubated for 2 h (37 °C, 5% CO_2_). No washing was performed after laminin coating.

OPCs were then seeded on the top of PDMS platforms or PLL pre-coated 13 mm diameter glass coverslips (50 µg/mL, 30 min, 37°C, and 5% CO_2_,) at a density of 600 viable cells/mm^2^. OPC seeding was performed using a 15 or 90 μL drop of cell suspension for micropillars and glass coverslips, respectively. Cell adhesion was promoted during 1 h – 1 h 30 min at 37 °C and 5% CO_2_. OPCs were maintained in OL SATO medium which was composed by DMEM medium supplemented with apo-transferrin (0.1 mg/mL, Merck T2036), bovine serum albumin (BSA) (0.1 mg/mL, NZYTech MB04601), putrescin (16 µg/mL, Sigma P7505), progesterone (60 ng/mL, Sigma P8783-), thyroxine (40 ng/mL, Merck T1775), sodium selenite (40 ng/mL, Sigma 214485-), triiodo-L-thyroxine (30 ng/mL, Merck T1775), human insulin (5 µg/mL, Sigma I9278), 0.5% heat-inactivated (30 min at 56 °C) FBS, and 1% (v/v) P/S. Cells were analysed at day 1 (D1), day 5 (D5), or day 7 (D7) of differentiation (DIFF). The overall purity of the OL culture was found to be above 70% for all substrates and micropillar diameters (Figure S3B).

For studies involving pharmacological microtubule stabilization analysis, cells were treated with paclitaxel (or Taxol®, Cytoskeleton Inc.) at 10, 25, 50, and 100 nM. Paclitaxel was added at D1 DIFF and incubated for 4, 8, 12, 24, and 48 h (37 °C, 5% CO_2_). Cells were then analyzed at D5 DIFF.

### Cellular metabolic activity (resazurin assay)

Metabolic activity was inferred by the measurement of resorufin fluorescence. Briefly, at D1, D5, and D7 DIFF, cells were incubated with resazurin solution (10% (v/v), in OL SATO medium) for 3 h (37 °C, 5% CO_2_). Supernatant medium was transferred to black 96-well plates and fluorescence measured at 530/590 nm in a multi-plate reader Synergy Mx (BioTeK® Instruments, GenX software).

### Cell viability (live-dead assay)

At D1, D5, and D7 DIFF OLs were incubated with a cell-permeant dye calcein-AM (1 µg/mL, λ_emission_ 517 nm/λ_excitation_ 484 nm, Promega) for 20 min (37 °C, 5% CO_2_). Cells were washed with phosphate buffered saline (PBS, pH 7.4) followed by a 5 min incubation at RT with propidium iodide solution (PI, 1 µg/mL, λ_emission_ 617 nm/λ_excitation_ 535 nm, Sigma-Aldrich), and Hoechst 33342 (10 µg/mL, λ_emission_ 497 nm/λ_excitation_ 361 nm, Molecular Probes). Cells were washed with PBS and immediately imaged under a widefield fluorescence microscope (37 °C, 5% CO_2_).

### Immunocytochemistry (ICC)

Cells were washed with pre-warmed (37 °C) PBS pH 7.4 and fixed with 4% (wt/v) microtubule-protecting paraformaldehyde solution (MP-PFA) for 20 min at RT. MP-PFA consists of 4% (wt/v) PFA (Merck Millipore), 65 mM PIPES (Millipore), 25 mM HEPES (Sigma-Aldrich), 10 mM EGTA (Sigma-Aldrich), 3 mM MgCl_2_ (Sigma-Aldrich) diluted in PBS pH 7.4 ^35^. OLs were further permeabilized and blocked with PBS containing 5% (v/v) normal donkey serum (NDS) (Sigma) and 0.3 % (v/v) Triton X-100 (Sigma). Primary antibodies were diluted in PBS containing 1% (v/v) NDS and 0.15% (v/v) Triton X-100 and incubated ON in a humid chamber at 4°C. The following primary antibodies were used: polyclonal rabbit anti-Oligodendrocyte Transcription Factor 2 (anti-Olig2) (1:400, Merck, AB9610) and monoclonal rat anti-myelin basic protein (anti-MBP) (1:100, Bio-Rad Laboratories, MCA409S). Secondary antibodies Alexa Fluor™ 647 anti-rabbit and 594 anti-rat (1:1000, ThermoFisher Scientific, A-11008 and A21209, respectively) and Alexa Fluor™ 488 Phalloidin (1:200, λ_emission_ 518 nm/λ_excitation_ 495 nm, ThermoFisherScientific, A12379) diluted in PBS containing 1% (v/v) NDS) and 0.15% (v/v) Triton X-100 were applied for 1 h at RT. Subsequently cells were treated for nuclear counterstaining at RT with Hoechst 33342 (1:1000, λ_emission_ 497 nm/λ_excitation_ 361 nm, Molecular Probes, H3570).

For studies involving pharmacological stabilization of the microtubules, monoclonal mouse anti-α tubulin (1:2000, Sigma-Aldrich, T5168) was used as a primary antibody and Alexa Fluor™ 647 anti-mouse as a secondary antibody (1:1000, ThermoFisher Scientific).

To evaluate cell culture purity, polyclonal rabbit anti-Ionized calcium binding adaptor molecule 1 (IBA-1, 1:500, Wako, 019-19741) was used to detect microglia, polyclonal rabbit anti-GFAP (1:1000, Abcam, AB7260) for astrocytes, rat anti-MBP for OLs, rabbit anti-Olig2 for OPCs/OLs, and monoclonal mouse anti-βIII tubulin (1:500, BioLegend, 801202) for neurons.

To understand the laminin distribution, micropillars were coated with laminin for 2 h and then OL SATO added for 10 min. The OL SATO was then replaced with pre-warmed PBS for 10 min and this step was repeated with fresh PBS for another 10 min. Micropillars were then fixed with 2% MP-PFA for 30 min. All the steps described above were performed at 37 °C, 5% CO_2_. Blocking was performed with 1% (wt/v) BSA and 4% (v/v) FBS. Primary antibody rabbit anti-laminin (1:200, Sigma, L939) was diluted in PBS and incubated overnight at 4 °C. Afterwards, micropillars were washed and incubated with secondary antibody (Alexa Fluor™ 488 anti-rabbit).

Following ICC, micropillars were mounted upside-down with Fluoromount aqueous mounting medium (Sigma, F4680, refractive index 1.40) in 35-mm ibidi dishes (ibidi 80136).

### Calcium imaging of oligodendrocytes

For calcium imaging studies, the single-wavelength calcium indicator Fluo-4 AM was used (Invitrogen F14201). At D5 and D7 DIFF, micropillars with OLs, cultured on #1.5 glass coverslip µ-Slide 8 well tissue culture treated and sterilized (ibidi), were incubated with Fluo-4 (4 μM) diluted in standard recording buffer for 30 min (37 °C, 5% CO_2_). The standard recording buffer was composed of 140 mM NaCl, 4 mM KCl, 2 mM CaCl_2_, 2 mM MgCl_2_, 10 mM HEPES, and 5 mM glucose diluted in type II distilled water (final pH 7.4). Cells were gently washed three times with recording buffer, incubated in the same solution for additional 15 min (37 °C, 5% CO_2_) and imaged immediately afterwards.

### Actin live imaging of oligodendrocytes

For studies involving actin live imaging of OLs, cells were incubated with Spy555-actin (SpiroChrome, 1:1000). For live imaging purposes, OL SATO was replaced by OL SATO diluted in FluoroBrite^TM^ DMEM (ThermoFisher) supplemented with GlutaMax and sodium pyruvate (1:100). Spy555-actin was incubated overnight at 37 °C, 5% CO_2_. Imaging was immediately performed without media washing or replacement in a TCS SP8 confocal laser scanning microscope.

### Confocal laser scanning microscopy

OLs on micropillars were imaged on a TCS SP8 confocal laser scanning microscope (Leica Microsystems). Images were acquired using a HC PL APO 20x /0.75 IMM/CORR CS2, with a resolution of 16 bits and in a sequential mode. A resonant frequency of 8000 Hz was applied with a bidirectional scan. An area of 465 x 465 μm with a pixel size of 323 nm (1024 x 1024, zoom factor 1.25) and a z-step size of 1.5 μm (z-size of 22.5 μm) were applied. 45 regions per PDMS platform were acquired. MBP labelled with Alexa Fluor 594 nm was excited with a HeNe 594 nm laser line with a laser power of 11% a gain of 31 and an offset of 0. Emission light was collected on a Leica HyD4 detector with a collection window of 604 – 642 nm. Olig2 labelled with Alexa Fluor 647 nm was excited with a HeNe 633 nm laser line with a laser power of 4% a gain of 11 and an offset of 0. Emission light was collected on a Leica HyD4 detector with a collection window of 643 – 726 nm. F-actin labelled with Alexa Fluor 488 Phalloidin was excited with an Argon laser with a laser power of 16% (set at 10% Argon laser power), a gain of 70 and an offset of 0. Emission light was collected on a Leica HyD3 detector with a collection window of 498 – 572 nm. Hoechst was excited with a 405 nm laser line with a laser power of 5.7%, a gain of 783 and an offset of −2. Emission light was collected on a Leica PMT detector with a collection window of 416 – 479 nm. Micropillar visualization was performed using transmitted light (gain 551, offset 0). The pinhole size was 56.6 μm, calculated at 1 airy unit (AU) for 580 nm emission.

For studies involving the live imaging of OLs labelled with Spy555-actin, timelapse images were acquired using a Fluotar VISIR 25x/0.95 WATER objective, with a resolution of 16 bits. A frequency of 400 Hz was applied with a unidirectional scan. An area of 155 x 155 μm with a pixel size of 152 nm (1024 x 1024, zoom factor 3) was applied. Spy555-actin was excited with a DPSS 561 nm laser line with a laser power of 0.6% (gain 100, offset −0.01). Emission light was collected on a HyD 3 detector with a collection window of 571 – 638 nm. Micropillar visualization was performed using transmitted light with a gain 527.6, offset 0. The pinhole size was 56.6 μm, calculated at 1 airy unit (AU) for 580 nm emission. Recordings were performed at 37 °C, 5% CO_2_ at 2.5 s per frame for a total period of 30 min.

For laminin 211 distribution studies, micropillars were imaged in a TCS SP5 confocal laser scanning microscope (Leica Microsystems) using a HCX PL APO CS 40.0×1.10 WATER UV objective, with a resolution of 16 bits. A frequency of 400 Hz was applied with a unidirectional scan and a line average of 3. An area of 194 x 194 μm with a pixel size of 190 nm (1024 x 1024, zoom factor 2) and a z-step size of 0.17 μm (z-size between 6 and 8 μm) was applied. Laminin was excited with an Argon laser (set at 20% Argon laser power) line with a laser power of 12% (gain 868.3, offset 0). Emission light was collected on a PMT2 detector with a collection window of 498nm - 584nm. Micropillar visualization was performed using transmitted light with a gain 377.3, offset 0. The pinhole size was 77.2 μm, calculated at 1 airy unit (AU) for 580 nm emission.

### Stimulated emission depletion microscopy (STED)

To evaluate myelin wrapping around micropillars, OLs at D5 and D7 DIFF were fixed and stained as previously described. Secondary antibodies consisted of 635 anti-rat STAR Red (Aberion, diluted 1:100). Samples were mounted in 90% (wt/v) glycerol medium (refractive index 1.44) and using a #1.5H μ-dish glass bottom 33 mm high (ibidi). Image acquisition was carried on a Leica Stellaris 8 STED Falcon microscope (single point scanning confocal equipped with a fully motorized inverted Leica DMI8 microscope, a white light laser (WLL), Falcon and STED modalities) from Leica Microsystems. Images were acquired using a HC PL APO CS2 93x/1.30 glycerol immersion objective equipped with a correction collar with a resolution of 12 bits. Images were recorded by frame scanning unidirectional at 400 Hz, with a line accumulation of 16, a pixel size of 20 nm, for an area size of 20.83 x 20.83 μm (1024 x 1024, zoom factor 6). For every micropillar imaged, four different planes were acquired (z size varied between 4 and 10 μm). A total of 20 micropillars were imaged per condition. MBP labelled with Abberior STAR 635P was excited with a 633 nm laser line with a laser power of 2%, and a gain of 10. Emission light was collected on a Leica HyDX2 detector (counting mode) with a collection window of 642 – 750 nm. The STED depletion was performed with a synchronized pulsed 775 nm (set at 85%) and a depletion laser at 50%. TauSTED was applied with the following parameters: Tau background: ON; Tau strength: 100; Denoise: 50. The pinhole size was 151.8 μm, calculated at 1 airy unit (AU) for 580 nm emission.

### Widefield fluorescence microscopy

Widefield fluorescence microscopy was used for live-dead assays, for studies on the pharmacological stabilization of OL microtubules, for calcium imaging, and to estimate the purity of the OLs on the PDMS platforms. For the live dead assay, OLs were imaged using a Leica DMI6000 widefield microscope coupled with a Hamamatsu-Flash4-CL-750935 (Hamamatsu, Japan) camera with temperature and CO_2_ controlled (37 °C, 5% CO_2_). Images were acquired using a HC FL PLAN 10x/0.25 DRY objective with a resolution of 16 bits, a pixel size of 500 nm, for an area size of 1024 x 1024 µm (2048 x 2048). Images were recorded using 100% camera light. 42 regions per PDMS platform (or glass coverslip) were acquired. Calcein-AM was recorded with the 527 nm emission wavelength filter (cube L5, 504 ms time of exposure), PI with the 645 nm emission wavelength filter (cube TX2, 1000 ms time of exposure), Hoechst with the 450 nm emission wavelength filter (cube AT2, 1000 ms time of exposure). PDMS platforms were imaged using the transmitted light (61 ms time of exposure).

For the pharmacological stabilization of OL microtubules, fixed OLs were imaged using a Timelapse Leica DMI6000 widefield microscope coupled with a Hamamatsu-Flash4-CL-850188 (Hamamatsu, Japan) camera. Images were acquired using a HCX PL FLUOTAR L 20x/0.40 CORR Ph1 DRY objective with a resolution of 16 bits, a pixel size of 650 nm, for an area size of 665 x 665 µm (1024 x 1024) and using 100% camera light. 42 regions per PDMS platform were acquired. MBP labelled with the Alexa Fluor 594 nm fluorophore was recorded with the 624 nm emission wavelength filter (cube TX2, 150 ms time of exposure). Olig2 labelled with the Alexa Fluor 488 fluorophore was recorded with the 527 nm emission wavelength filter (cube L5, 200 ms time of exposure). α-tubulin labelled with the Alexa Fluor 647 was recorded with the 700 nm emission wavelength filter (cube Y5, 150 ms time of exposure). Hoechst was recorded with the 470 nm emission wavelength filter (cube AT, 30 ms time of exposure). PDMS platforms were imaged using the transmitted light (15 ms time of exposure).

For calcium imaging experiments, OLs were imaged using the same microscope, objective, resolution, pixel size and area size as for studies on pharmacological stabilization of OL microtubules. To detect spontaneous changes in intracellular calcium concentration, Fluo-4 fluorescence was recorded with the 527 nm emission wavelength filter (cube L5, 100 ms time of exposure) using 20% of the camera light. Recordings were performed at RT at 1.2 s per frame for a total period of 5 min.

To estimate the purity of the OLs on the different PDMS platforms, fixed and stained cells were imaged using the INCell Analyzer 2000. For glass coverslips imaging, 25 equally spaced fields were acquired in a 2D mode. For PDMS surface and micropillars imaging, nine fields (3×3 grid of juxtaposed fields) were acquired per diameter, in a 2.5D mode (3 μm depth).

### Image segmentation and analysis

To quantify spontaneous calcium transients, each time-lapse image was processed to generate a ΔF/F_0_ sequence. Using ImageJ (version 1.52u 17 March 2020, https://imagej.nih.gov/ij/notes.html) ^36^, a maximum z projection image was created for all timelapse sequences and then regions of interest (ROI) were defined based on manual thresholding (adjusted for every image analyzed). A watershed filter was applied to separate cell clusters and then cells were segmented using the “Analyze particles” tool, applying an area filter (minimum 20 μm^2^ to avoid inclusion of cell debris). Afterwards, every ROI was individually verified. Additionally, if a cell was not included in the first ROIs, manually drawn ROIs were applied. For every image analyzed, a background ROI was also manually included. After applying the defined ROIs to the timelapse images, the “Time Series Analyzer” plugin from ImageJ was initialized and the raw fluorescent intensity values for all the ROIs were extracted. Further analyses were carried out in Excel and Python. For each cell, raw fluorescence intensity was normalized with respect to the mean raw intensity values of the first 10 frames where no spontaneous activity occurred (baseline fluorescence, F_0_), after background subtraction. For every ROIs, the change in fluorescence intensity with respect to the baseline, ΔF 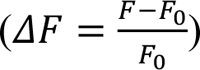, was estimated and individually plotted. We scored calcium events as increases of ΔF/F_0_ equal to or more than 40% of the baseline. Cells that did not have an increase above this level were considered “inactive”; ROIs where bleaching occurred were excluded and ROIs with linear increase on the fluorescence values were also excluded from the analysis. The percentage of active cells, the peak amplitude and the number of calcium events (frequency) were then estimated for every single cell.

For all image analysis performed regarding OL differentiation and myelination, images were firstly converted from a. lif to a .tiff format using the ImageJ (version 1.52u 17 March 2020, https://imagej.nih.gov/ij/notes.html) ^36^. To assess the Olig2 and MBP cell number, area of MBP and OL differentiation stage, channels were split, and z-stack maximum projection images were used. To evaluate the number of myelinated micropillars the best z-plane for each image was chosen. Probability maps for all channels were then generated using ilastik (version 1.3.3, https://www.ilastik. org/) ^37^. Further image segmentation and analysis was performed using CellProfiler (version 3.1.9, https://cellprofiler.org) ^38^. Briefly, nuclei were identified from Hoechst images by detection of objects with sizes ranging from 7 to 40-pixel units using the Otsu threshold method (threshold smoothing scale 1.3488, threshold correction factor 0.4) and clumped objects were distinguished by shape. To estimate the number of MBP positive cells, secondary objects were identified by a 5-pixel expansion of the nuclei objects, and the intensity of the MBP was estimated in the expansion of the nuclei area. Olig2 image intensity was measured in the nuclei objects. The MBP objects were detected from the MBP channel images using a manual threshold method (0.35 and threshold smoothing scale 1.3488). The overall MBP image intensity as well as MBP occupied area was estimated. Property files were then transferred to CellProfiler Analyst (version 3.0.4) to categorize cells as positive or negative for MBP and Olig2. The random forest classifier algorithm for object classification was chosen and classification accuracies above 85% were accepted. For estimation of OL culture purity as well as for calculating the number of viable/dead cells, a similar protocol was followed. Cells were categorized either in GFAP, IBA-1, βIII tubulin, Olig2, and MBP positive or Calcein and PI positive.

To quantify the micropillars wrapping by OLs, micropillars were detected using a manual threshold technique (0.25, threshold smoothing scale 1.3488) of objects in the range of 1-to-20-pixel units of size. Depending on the micropillar diameter, different filters were applied to discard debris erroneously identified as micropillars objects. Based on the best myelinated z image, the “MBP in pillars” objects were identified by a manual threshold (0.3, threshold smoothing scale 1.3488) and masked in a 2-pixel expansion of micropillars objects (to consider only the signal corresponding to the myelin rings). The total number of micropillars as well as the presence/absence of myelin around the micropillars was estimated along with the MBP intensity in the micropillars. Additionally, to calculate the total area of the MBP around the micropillars, a 20-pixel size expansion from the micropillars object was created.

For classification of OLs differentiation stage, the collar occupancy method developed and published by Bouçanova et *al* was adapted ^22^. Briefly, a collar object was created by adding a 12-pixel unit expansion to the nuclei. The F-actin objects (detected via a manual threshold of 0.6, threshold smoothing scale 1.3488) were then masked by the collar objects and the occupancy area of the F-actin estimated. Only cells positive for the Olig2 staining (nuclei containing a mean intensity of the Olig2 probabilities map image above 0.150) and single cells (generally with nuclei area between 50 and 400 and collar area between 700- and 1200-pixel units) were considered for the analysis. Collar occupancy areas were defined for each differentiation stage: stage I area below 30-pixel units, stage II area between 31- and 53-pixel units, stage III area between 54- and 66-pixel units, and stage IV area above 67-pixel units.

For OL myelin thickness and *g*-ratio measurements, the length of the MBP staining across the micropillar was manually measured using ImageJ (version 1.52u 17 March 2020, https://imagej.nih.gov/ij/notes.html) ^36^. A total of six measurements per z-plane were performed and one single measurement of the real value of the micropillar was inferred from the middle z-plane.

To measure the motion of the cells around the pillars Image Velocimetry (PIV) was implemented. PIV was analyzed using the MatPIV toolbox in regions containing cells previously delineated using MATLAB code described by us ^39^. PIV measures a velocity field from time-lapse images by dividing each frame into multiple regions and calculating by how much and in what direction that region has shifted in the next frame. The calculations were conducted over four iterations, with the initial two iterations using 64 pixel × 64 pixel regions, followed by two iterations using 32 pixel × 32 pixel regions. PIV determines the most likely shift of a region from measurements of correlations between subsequent frames. The resulting PIV vector field was then translated into radial and tangential motion relative to the nearest pillar’s pixel locations.

### RNA extraction, cDNA synthesis and quantitative real-time polymerase chain reaction (qRT-PCR)

Total RNA from OPCs was extracted using the Quick-RNA MiniPrep kit (Zymo Research R1050) according to the manufacturer’s recommendations. Eight micropillar platforms from the same condition were pooled together. RNA concentration and purity was estimated using the NanoDrop 1000 Spectrophotometer (ThermoFisher Scientific) and its integrity was assessed using Experion^TM^ RNA StdSens Analysis Kit (Biorad). Afterwards, cDNA was synthesized from 100 ng of RNA using the NZY First-Strand cDNA kit (NZY, MB12501) following the manufacturer’s protocols. Quantitative real-time PCR was performed on CFX 384 (Bio-rad) in triplicates using the iTaq Universal SYBR Green Supermix (Bio-rad) composed by 0.25 μL for each primer (final concentration 25 nM), 5 μL of iTaq, 1 μL of cDNA, and 3.5 μL of RNAse free water. The melting temperature was optimized for 55 °C for all the primers tested. Primer gene sequences were designed using the NCBI primer tool and the Beacon designer software and can be found in Table S3. The amplification efficiencies were tested for all primer pairs and were only used when efficiency was near 100%. To verify the specificity of the amplification and absence of primer dimer formation, corresponding melting curves were performed and analyzed immediately after the amplification protocol. Non-specific products were not found in any case. *Oaz1* was used as endogenous control to normalize the expression levels of genes of interest and the relative mRNA expression levels were calculated using the delta C_T_ (2^-ΔΔCT^) method ^40^.

### Statistical analysis

Statistical analysis was performed using GraphPad Prism (version 7.00, www.graphpad.com). For all datasets, tests for identification of outliers and normality were performed. To identify and remove likely outliers, the Robust Regression and Outlier Removal (ROUT) method with Q=10% was used. Following this, Gaussian distributions were tested using the Shapiro-Wilk normality test (α=0.05). When all datasets followed a normal distribution, statistical differences between groups were calculated based on one-way (influence of one factor) or two-way ANOVA (influence of two factors), followed by Dunnet’s or Tukey’s multiple comparisons test, respectively, for multiple comparisons. In case one dataset did not follow a normal distribution, non-parametric tests were performed. Statistical differences between groups were calculated based on Kruskal-Wallis analysis, followed by Dunn’s multiple comparisons test for multiple comparisons. A p-value below 0.05 was considered statistically significant and data are shown as mean ± standard deviation (SD).

## Acknowledgements

The authors acknowledge the support of the i3S Scientific Platforms Bioimaging, Advanced Light Microscopy (ALM) and Histology and Electron Microscopy (HEMS), members of the PPBI (PPBI–POCI-01-0145-FEDER-022122). The authors thank Ana Leite Oliveira and Fernanda Isabel Ferreira from Universidade Católica Portuguesa for the kind help with PDMS refractive index analyses. The authors thank Centro de Materiais da Universidade do Porto (CEMUP) for SEM and AFM analyses and Manuela Brás from i3S for the kind help in analyzing the AFM data. The authors acknowledge Joana Bravo and Miguel Aroso from i3S for kindly sharing spared rat primary oligodendrocytes cultures whenever available. This work was supported by FCT – Portuguese Foundation for Science and Technology – UT Austin Portugal Program (UTAPEXPL/NTec/0057/2017) and Air Force Office of Scientific Research, USA (Awards FA9550-20-1-0417 and FA9550-23-1-0599). EDC, MRGM and SCG acknowledge the support by FCT – Portuguese Foundation for Science and Technology their fellowships (SFRH/BD/140363/2018; 2022.13353.BD; and SFRH/BPD/122920/2016 respectively). GA acknowledges the support by MCSA ASTROTECH 956325-ASTROTECH-H2020-MSCA-ITN-2020. EDC acknowledges COST Action CA16122 (BIONECA) for supporting a visit to Prof. Eduardo Mendes lab at TU Delft. APP acknowledges the MOBILIsE Project, which has received funding from the European Union’s Horizon 2020 research and innovation program under grant agreement no.951723.

## Author contributions

Conceptualization: EDC, APP; Methodology: EDC, MRGM, HPF, GA, HH, RR, BC; Investigation: EDC, MRGM, HPF, GA, HH, RR, MM, BC, SG, KAYPS, SLR, SC, SCG; Visualization: EDC; Funding acquisition: APP; Supervision: SC, WL, SCG, EM, APP; Writing—original draft: EDC; Writing—review & editing: MRGM, GA, HH, BC, MGL, SCG, APP.

## Notes

### Competing Interest Statement

The authors have declared no competing interest.

