## Supplemental Figures and Tables for "Engineered micropillars to unveil oligodendrocyte responses to physical cues"

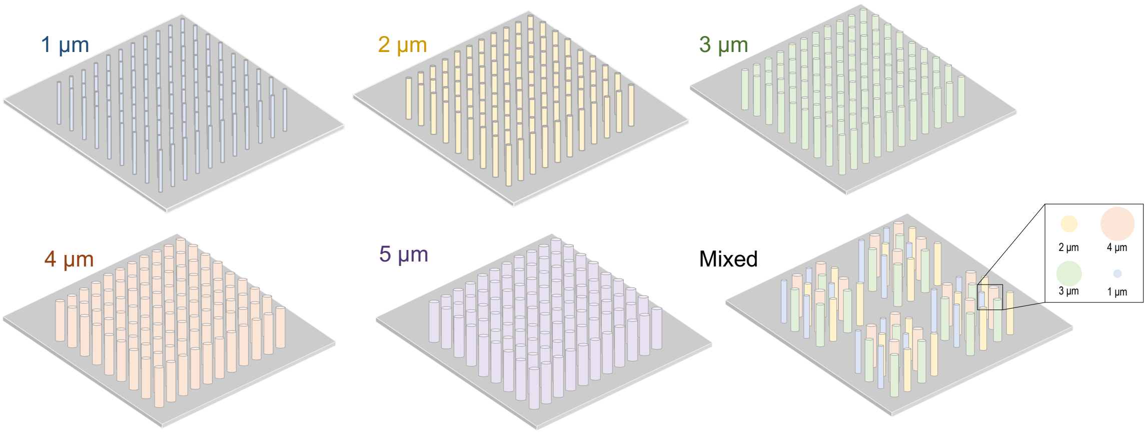


**Figure S1. Representation of the micropillars platforms used in the present study.** Micropillars were projected to have diameters ranging from 1 to 5 µm, a heigh of 10 µm and interspaced 30 µm. For live cell imaging purposes, a mixed micropillar platform consisting of 1 to 4 µm diameter microstructures was designed.

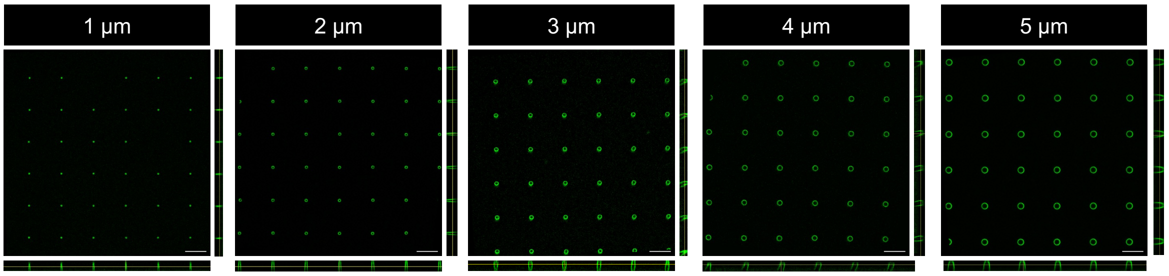


**Figure S2. Laminin distribution along PDMS micropillars.** Laminin 211 coating (in green) distribution on micropillars of 1, 2, 3, 4 and 5 µm (confocal representative images). Orthogonal views are represented for each image. Scale bar indicates 20 µm.

A

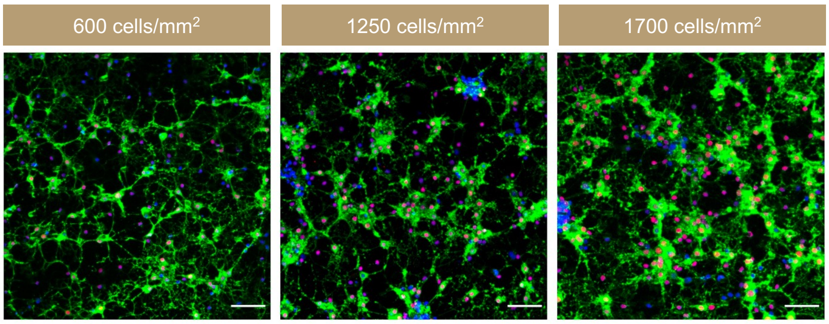


B

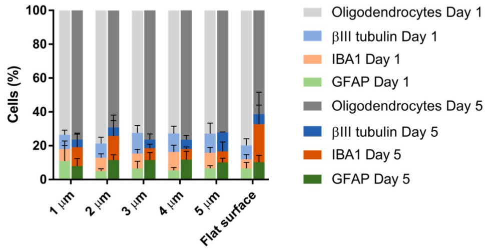


**Figure S3. Oligodendrocyte (OL) culture conditions optimization. (A)** Representative images of the cell density optimization. Three different cell densities were tested (600, 1250, and 1700 cells/mm$^2$) and the lowest one was chosen for further studies, since it provided sufficient free area for OL branching and process extension. Mixed micropillars of rigid PDMS were used in these experiments. Green: myelin basic protein (MBP), blue: Hoechst, red: Olig2. Scale bar indicates 50 µm. **(B)** Estimation of OPC purity from cultures seeded on PDMS platforms at day 1 and day 5 of differentiation (DIFF). Percentage of oligodendroglial cells was calculated as the difference between 100% and the sum of average values for GFAP+, IBA1+ and βIII tubulin+ cells. Graphs present mean values with standard deviation (SD) bars. No SD bars for the oligodendroglial fraction. OPC purity was estimated to be approximately 75% at D1 DIFF, evolving to 72% at D5 DIFF.

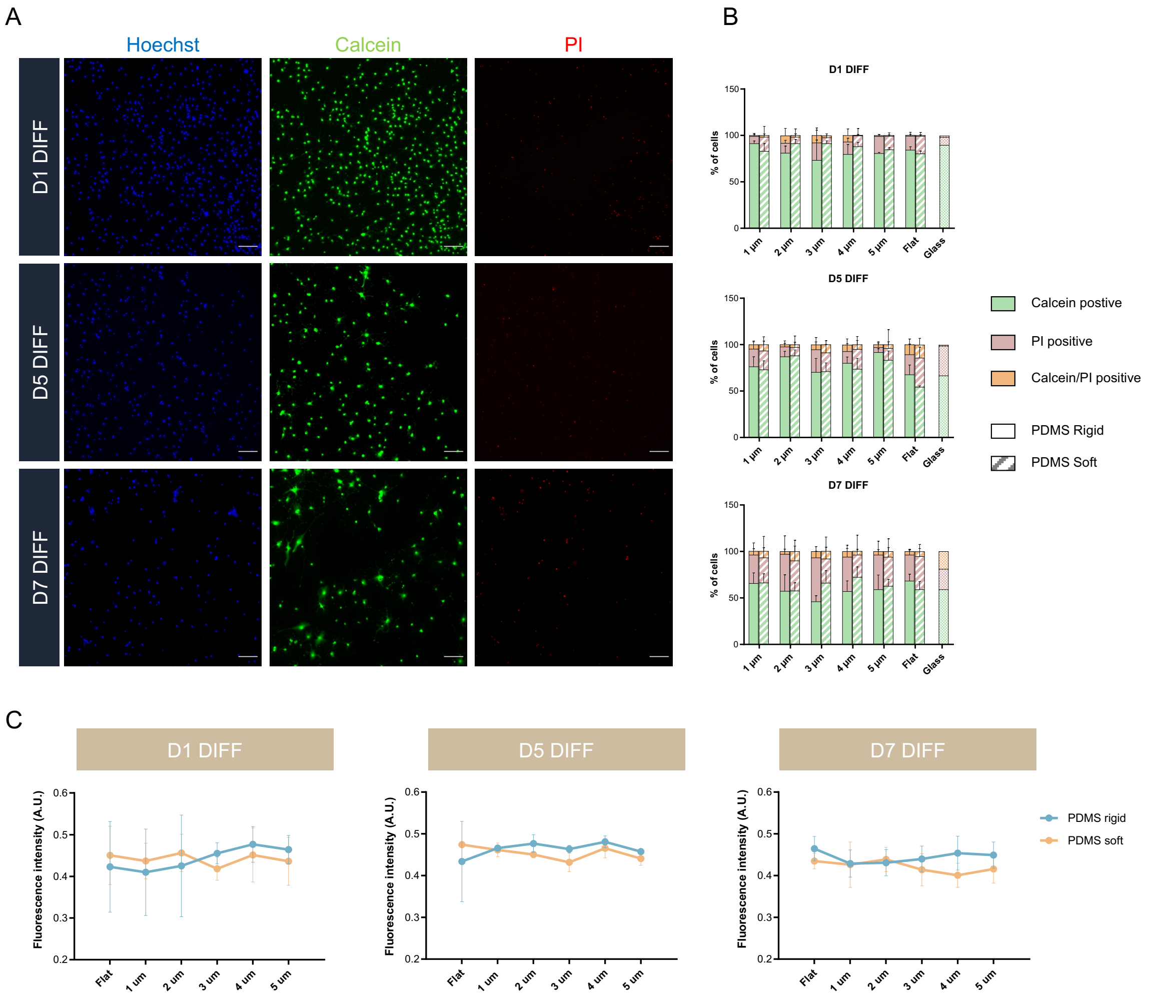


**Figure S4. Oligodendrocyte (OL) survival and metabolic activity when cultured on micropillars. (A)** Representative images of OL viability on micropillars at day 1 (D1), D5, and D7 of cell differentiation (DIFF). Images show OLs on 4 µm rigid microstructures. OLs were stained with Calcein-AM (green, showing the live cells), propidium iodide (PI, red, representing the death cells) and Hoechst (blue). Scale bar indicates 50 µm. **(B)** Quantification of the number of Calcein and PI positive cells for all micropillar structures tested. OLs growing on glass coverslips were used as quality control for the culture viability. Results show mean ± SD, n=3 independent experiments. Statistical analysis was performed using Two-way ANOVA, $^*p < 0.05$. **(C)** Metabolic activity of OLs on micropillars and flat PDMS structures at D1, D5, and D7 DIFF. Metabolic activity was inferred by means of the resazurin assay. Results show mean ± SD, n=3 independent experiments, two replicates per experiment. Statistical analysis was performed using Two-way ANOVA.

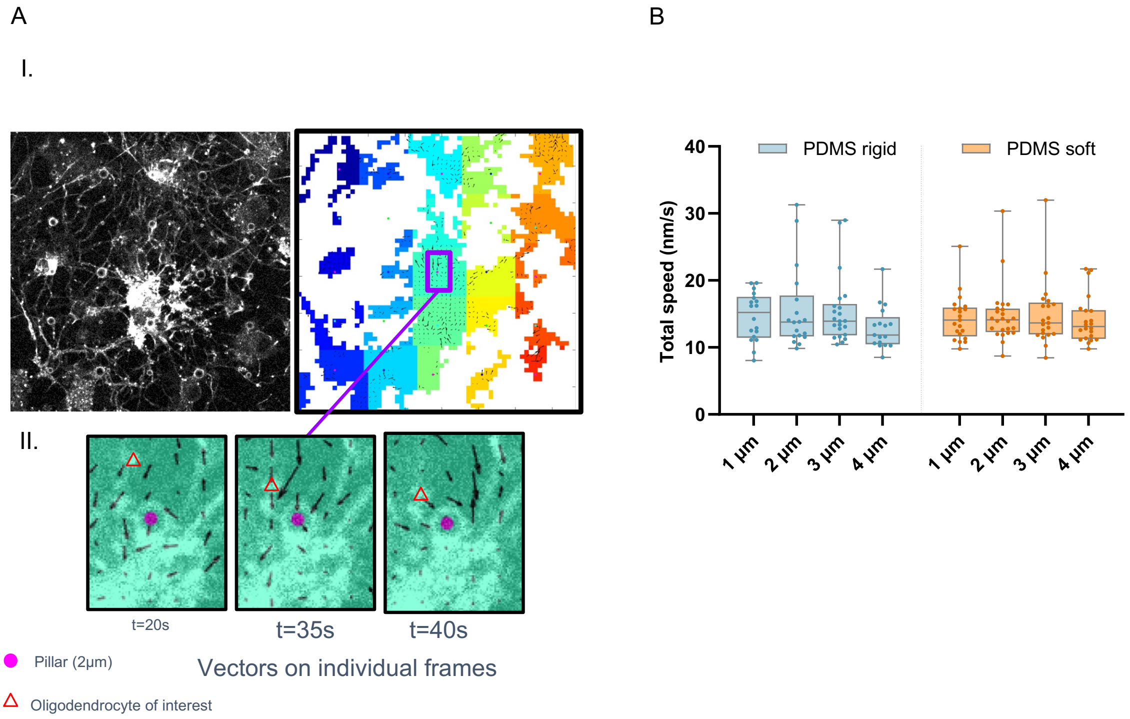


**Figure S5. Oligodendrocyte particle image velocimetry (PIV) analysis at day 5 of differentiation (DIFF). (A)** Representative image of the PIV vector field (I) and amplified region of PIV vectors overlaid on top of original Spy555-Actin fluorescence image. The red triangle marks an oligodendrocyte (OL) moving closer to a 2 µm micropillar over time. **(B)** Mean comparison of the total speed of OLs in the region around the different micropillars. Statistical analysis was performed using Two-way ANOVA.

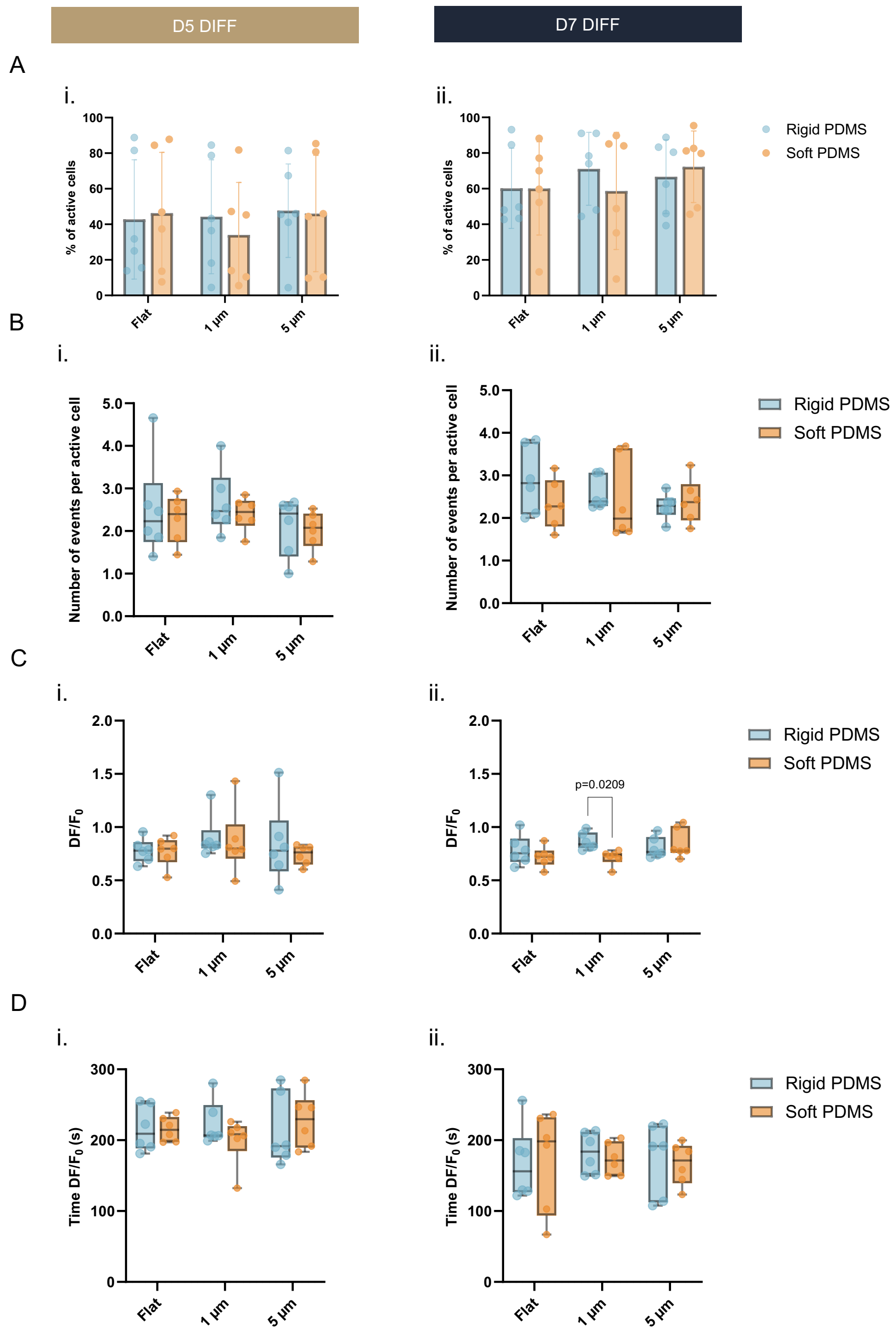


**Figure S6. Oligodendrocyte (OL) calcium signaling events on micropillars at day 5 and 7 of differentiation (DIFF). (A)** Quantification of the number of active cells on flat, 1 and 5 µm rigid and soft surfaces at D5 and D7 DIFF, respectively. **(B)** Frequency of calcium events quantified for every cell analyzed at D5 and D7 DIFF, respectively. **(C)** Peak amplitude of the calcium event at D5 and D7 DIFF. **(D)** Time of the maximum peak amplitude at D5 and D7 DIFF. n = 6 images analyzed per condition (from 3 independent experiments) with > 120 cells per image. Two-way ANOVA.

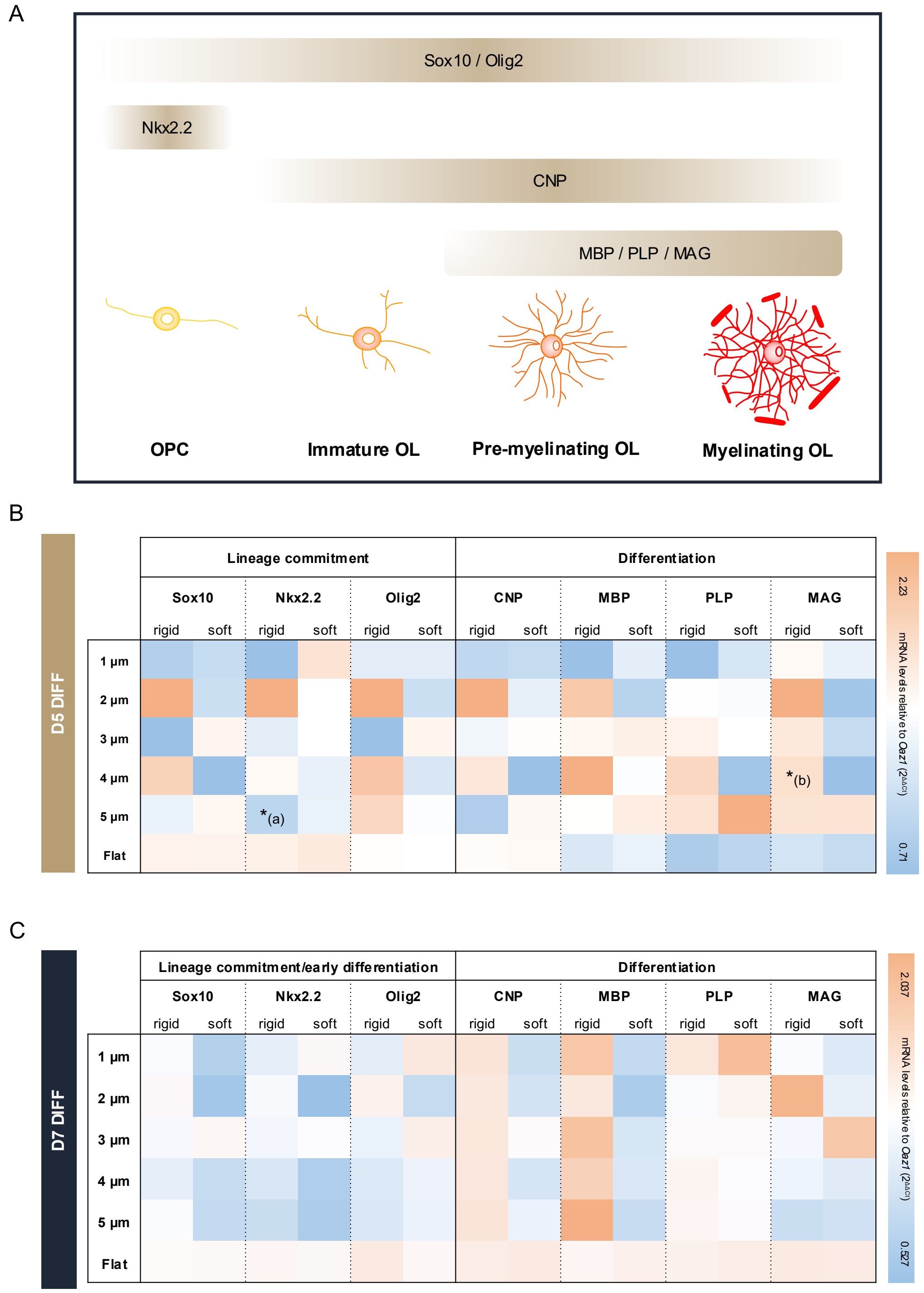


**Figure S7. Oligodendrocyte (OL) gene expression from cells growing on different PDMS structures at D5 and D7 DIFF. (A)** Representative scheme of the oligodendrocyte precursor cell (OPC) and OL characteristic markers. Scheme based on Gregath et al (2018) and Kamen et al (2022). **(B)** and **(C)** mRNA levels of OLs on PDMS structures at D5 and D7 DIFF, respectively. Results indicate mean values from three independent experiments (eight micropillar/flat structures pooled together). Lineage commitment/early differentiation (Sox10, Nkx2.2, and Olig2) as well as differentiation (CNP, MBP, PLP, and MAG) markers were analyzed. Levels were normalized to Oaz1 housekeeping gene and to respective flat surfaces (2-ΔΔCt, n>3 independent experiments, 8 micropillar platforms pooled together). Statistical analysis was performed using Two-way ANOVA. a) represents statistically differences between 5 µm and flat rigid surfaces and b) represents statistically differences between 4 µm rigid and soft micropillars. * $p \leq 0.05$.

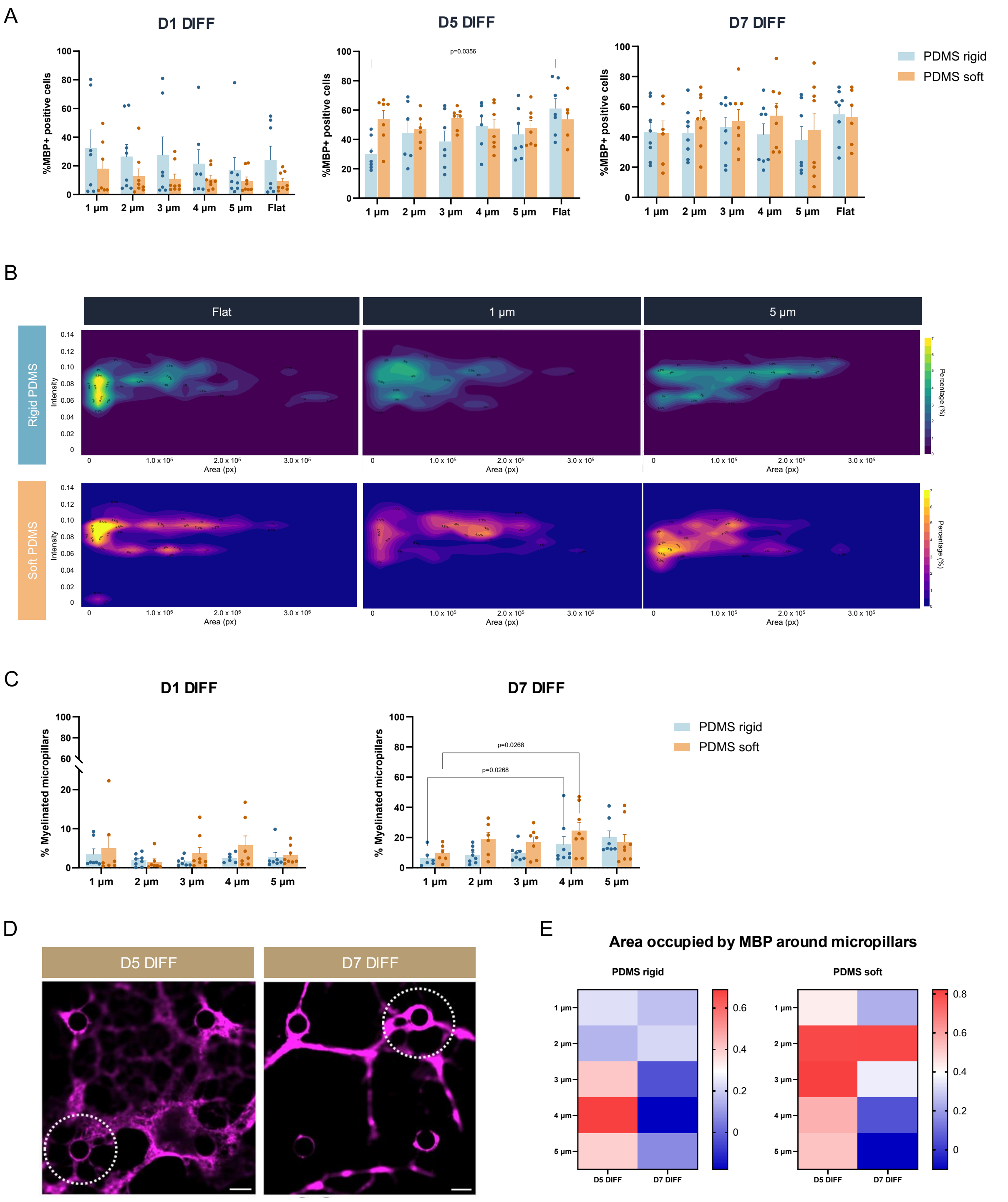


**Figure S8. Effect of micropillar diameter and stiffness on oligodendrocyte (OL) differentiation and wrapping. (A)** Percentage of myelin basic protein (MBP) cells throughout cell culture time. Results show ± SD, n > 5 micropillars platforms from 3-4 independent experiments. Statistical analysis was performed by Two-way ANOVA. **(B)** Density distribution graphs of the MBP area as a function of its mean intensity. For all the samples the mean intensity values were normalized for flat PDMS rigid surfaces. n>250 images analyzed per condition (from 4 independent experiments). **(C)** Percentage of wrapped micropillars by OLs on micropillars of different diameters and stiffness at D1 and D7 DIFF. Results show mean ± SD, n>6 micropillar arrays analyzed from four different independent experiments.

Statistical analysis was performed by Two-way ANOVA. **(D)** Representative images of OL wrapping around micropillars at day 5 and 7 of differentiation. White dotted circles note the region considered for the quantification of area occupied by the myelin basic protein (MBP). Scale bar indicates 5 μm. **(E)** Quantification of the MBP area around micropillars. Results show mean values. ($n > 3$ independent experiments, two replicates per experiment).

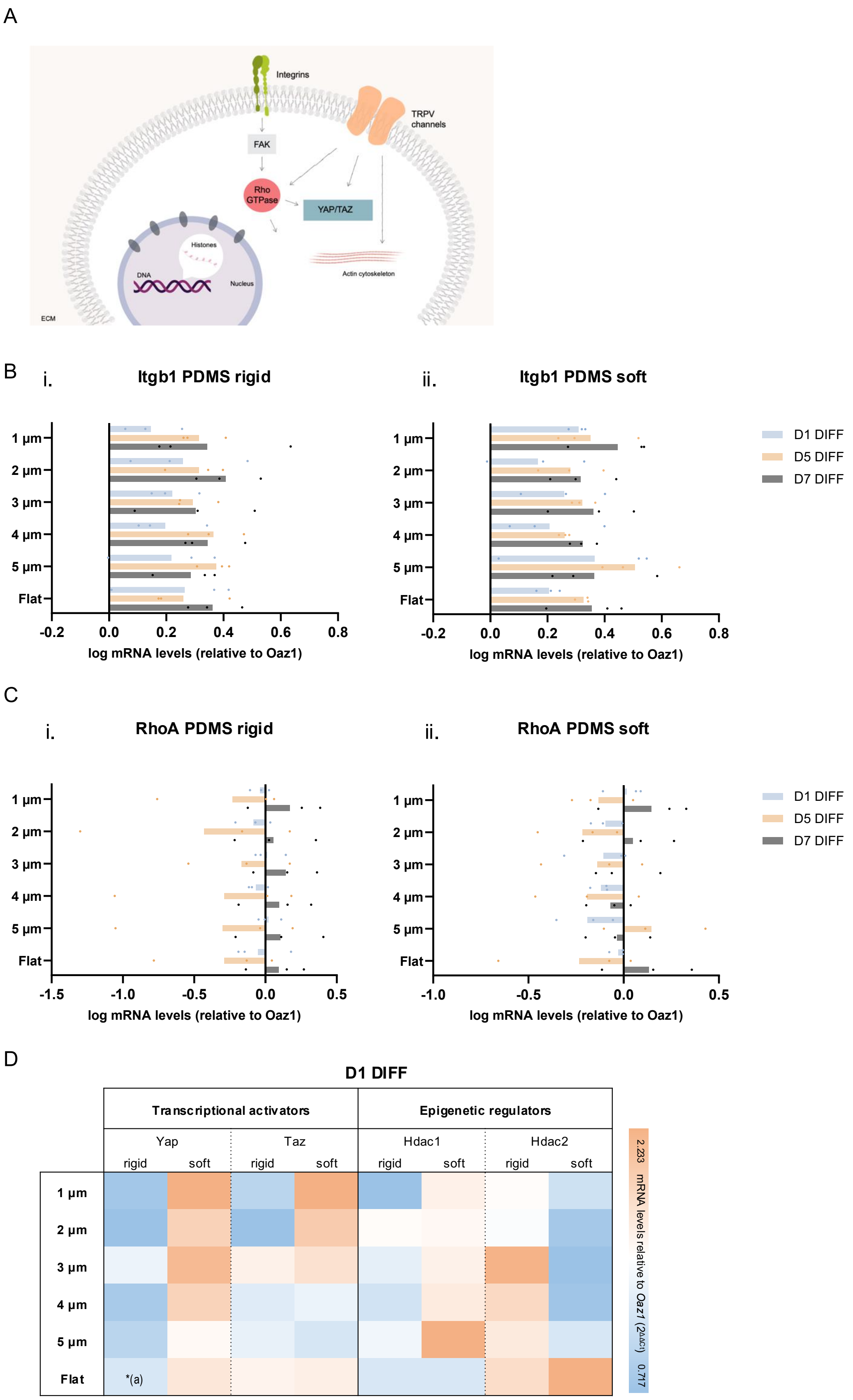


**Figure S9. Cytoskeleton related genes expressed on oligodendrocytes over time of culturing. (A)** Scheme of the mechanosensing key regulators studied in the context of this work. **(B)** and **(C)** mRNA levels of integrin beta 1 receptor and RhoA, respectively. Gene expression levels were normalized to

*Oaz1* housekeeping gene ($2^{-\Delta\Delta Ct}$, n=3 independent experiments, 8 micropillar platforms pooled together). Statistical analysis was performed using Two-way ANOVA. **(D)** mRNA expression levels of mechanosensing genes (*Yap*, *Taz*, *Hdac1* and *Hdac2*) of OLs cultured on micropillars at day 1 of differentiation. Results show mean values, n>3 independent experiments (eight micropillar platforms pooled together for each condition per independent experiment). Statistical analysis was performed using Two-way ANOVA, p*<0.05. (a) significantly different between 1 μm and flat rigid.

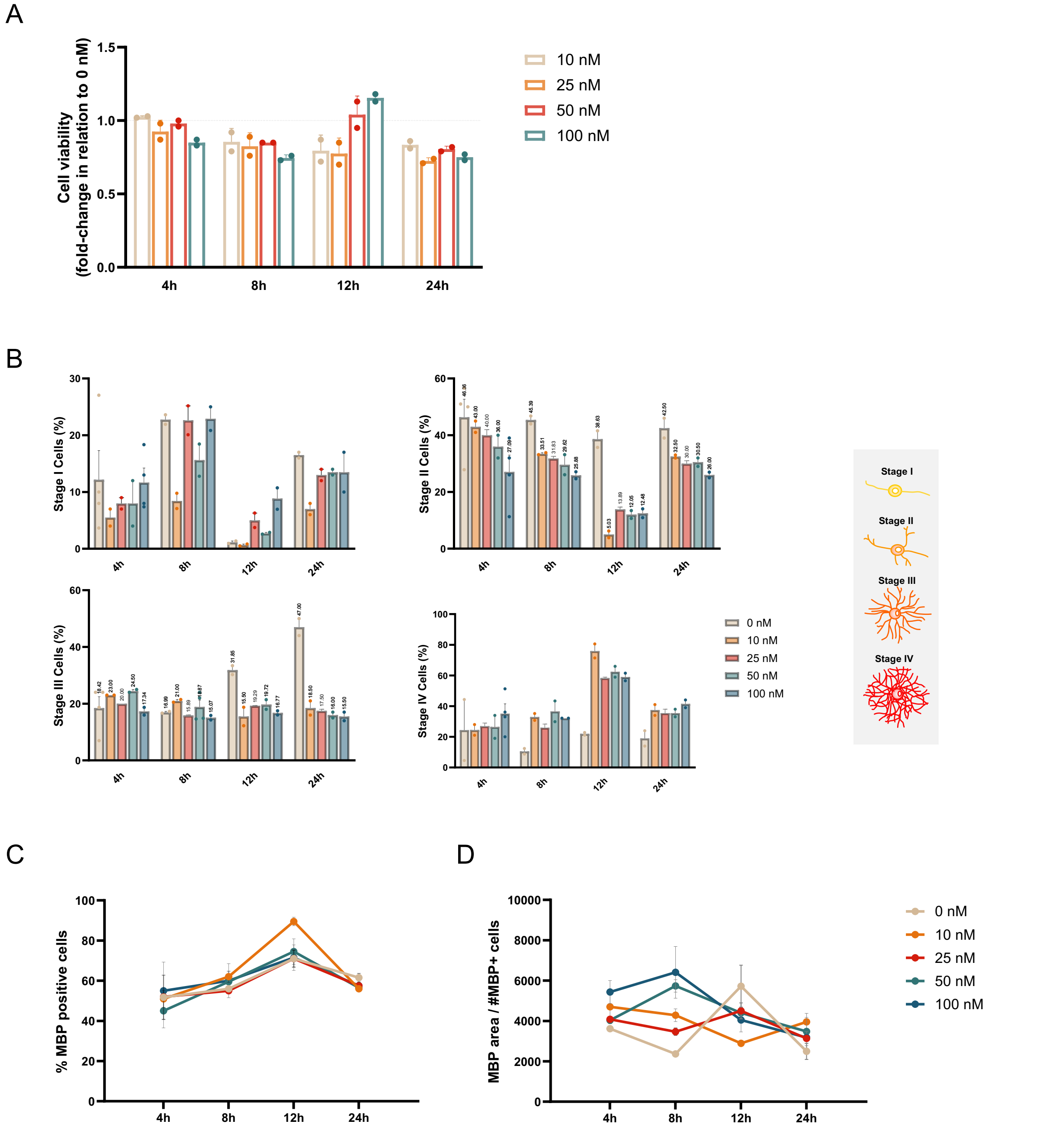


**Figure S10. The Effect of Paclitaxel (Taxol) on oligodendrocyte (OL) cultures.** A screening of different concentrations (10, 25, 50 and 100 nM) at different exposure times (4, 8, 12, and 24 h) was performed. Taxol was added to the cultures 18 h after cell seeding and the analysis was performed at day 5 of differentiation (DIFF). **(A)** Cell viability of OLs cultured in the presence of Taxol on glass coverslips. Results show mean ± SD, n = 2 independent experiments (2 replicates per experiment). **(B)** Quantification of OL branching ability OL. Olig2+ cells were grouped into four different stages where the stage I represents the bipolar OPCs and the stage IV the highly branched OLs. Results represent mean ± standard deviation (SD), n = 2 independent experiments (2 replicates per experiment). **(C)** and **(D)** MBP positive and MBP occupied area after treatment with Taxol. Results represent mean ± standard deviation (SD), n = 2 independent experiments (2 replicates per experiment).

**Table S2. In vitro platforms reported in the open literature used to mimic oligodendrocyte myelination.**

| Reference | 2.5D platform? | Tunable stiffness? | Diameter compatible with CNS diameters? | Mixed diameters? | Myelination followed by live-imaging? | Amenable for high-throughput? |
|---|---|---|---|---|---|---|
| Jagielska et al, 2012 [9] | No (2D scaffold) | Yes | - | - | - | Yes |
| Urbanski et al, 2016 [6] | No (2D scaffold) | Yes | - | - | - | Yes |
| Segel et al, 2019 [8] | No (2D scaffold) | Yes | - | - | - | Yes |
| Howe et al, 2006 [21] | Yes (microfibers) | No | Yes | No | No | No |
| Lee et al, 2012 [12] | Yes (microfibers) | No | Yes | No | No | Yes |
| Bechler et al, 2015 [13] | Yes (microfibers) | No | Yes | No | No | Yes |
| Ong et al, 2020 [20] | Yes (microfibers) | Yes | Yes | No | No | Yes |
| Mei et al, 2014 [14] | Yes (micropillars) | No | No | No | No | Yes |
| Espinosa-Hoyos et al, 2018 [15] | Yes (micropillars) | Yes | No | No | No | Yes |
| Yang et al, 2025 [16] | Yes (micropillars) | Yes | Yes | No | No | Yes |
| **PDMS micropillars** | **Yes (micropillars)** | **Yes** | **Yes** | **Yes** | **Yes** | **Yes** |

**Table S2. Measurements of diameter at mid-length, base, and top of the micropillars, height (from lateral view SEM images) and distance between micropillars (from top view SEM images), for poly(dimethylsiloxane) (PDMS) micropillars.** Presented values correspond to an average of three measurements. Results show mean ± standard deviation (SD). The distance between micropillars was measured from micropillars' wall to wall.

| Micropillar | Diameter at mid-length (µm) | Diameter at base (µm) | Diameter at top (µm) | Height (µm) | Distance between micropillars (µm) |
|---|---|---|---|---|---|
| 1 µm rigid | 1.318 ± 0.011 | 1.541 ± 0.067 | 1.207 ± 0.016 | 6.559 ± 0.181 | 28.491 ± 0.586 |
| 2 µm rigid | 2.431 ± 0.019 | 3.116 ± 0.051 | 2.317 ± 0.038 | 9.231 ± 0.019 | 28.463 ± 0.472 |
| 3 µm rigid | 3.574 ± 0.014 | 4.478 ± 0.066 | 3.234 ± 0.050 | 10.664 ± 0.029 | 28.330 ± 0.614 |
| 4 µm rigid | 4.775 ± 0.011 | 5.450 ± 0.132 | 4.397 ± 0.011 | 11.880 ± 0.058 | 27.912 ± 0.752 |
| 5 µm rigid | 5.871 ± 0.087 | 6.900 ± 0.125 | 5.292 ± 0.029 | 12.786 ± 0.087 | 28.018 ± 0.288 |
| 1 µm soft | 1.247 ± 0.023 | 1.603 ± 0.066 | 1.196 ± 0.022 | 5.752 ± 0.022 | 25.801 ± 0.449 |
| 2 µm soft | 2.150 ± 0.022 | 2.545 ± 0.058 | 1.985 ± 0.038 | 7.418 ± 0.044 | 25.702 ± 0.316 |
| 3 µm soft | 3.066 ± 0.058 | 3.525 ± 0.059 | 2.812 ± 0.022 | 8.296 ± 0.045 | 25.536 ± 0.615 |
| 4 µm soft | 4.071 ± 0.044 | 5.065 ± 0.045 | 3.776 ± 0.059 | 9.428 ± 0.067 | 25.470 ± 0.289 |
| 5 µm soft | 5.230 ± 0.065 | 6.108 ± 0.115 | 4.722 ± 0.080 | 10.599 ± 0.023 | 25.501 ± 0.249 |

**Table S3 Primer sequences used for oligodendrocyte gene expression studies.** Forward and reverse sequences are shown.

| Primer name | Forward sequence (5'-3') | Reverse sequence (5'-3') |
|---|---|---|
| **MBP** | TGTCACAATGTTCTTGAAGAA | GCTCCCTGCCCCAGAAGT |
| **MAG** | TTATTGTTATCCTGGTCTCC | ACTGGCACTTGTCATTAG |
| **PLP** | GCCAGAATGTATGGTGTTCT | TTAAGGACGGCAAAGTTGTA |
| **CNP** | GCTTCGACACTTCATTTCTG | GTCTCTTGCCAAAATAGCTG |
| **Nkx2.2** | TGCCCCTTAAGAGTCCTTT | TCCGTGCAGGGAGTATT |
| **Sox10** | AAGCTCTGGAGGTTGCTG | CGAGGTTGGTACTTGTAGTC |
| **Olig2** | TGGGTGTCAGAAACACTTAG | CGTTAGGAAACCACAAATCG |
| **Yap1** | TGGTGAGAGGTAGCAGAG | GCAAGATGAGAGCGAAGT |
| **Taz1** | AAGGAAGTGCTGTATGAG | AAGACTGGTGGTTAGAGA |
| **Hdac1** | GAGCAAGATGGCGCA | GCTTTGTGAGGACGATAGA |
| **Hdac2** | AAATTCCCAATGAGTTGCCA | TGTGGTAACATTCGCAGATT |
| **Hdac3** | TCTTTCCTGGAACAGGTGATA | ATCAATGCCATCCCGTAAG |
| **TRPV1** | GAATTGAGATGTGTATCCA | ATGTGTCCAAGTAGAGAT |
| **TRPV4** | CATTAACGAACTGCTGAG | ACACAGATAGGAGACAAC |
| **RhoA** | TGCTTGCTCATAGTCTTC | TTCAATATCTGCCACATAGT |
| **Itgb1** | ACCTCTAATCTCTAATGTGTCT | CAGGACGGCTTACAGTAT |
| **Oaz1** | TGCAGCAGCGAGAGTTCTAGG | CCGGACCCAGGTTACTACAG |

**Movie S1.**

Timelapse video of oligodendrocytes cultured on micropillars for 48h (from day 0 to day 2 of differentiation).

**Movie S2.**

Representative timelapse video of oligodendrocytes labelled with Spy555-Actin at D5 DIFF. Scale bar represents 25 µm.

**Movie S3.**

Representative timelapse video of calcium events on 1 µm **(A)**, 5 µm **(B)** PDMS rigid micropillars, and flat rigid surfaces **(C)** at day 5 of differentiation. Scale bar represents 25 µm.
